# PERIOD phosphorylation leads to feedback inhibition of CK1 activity to control circadian period

**DOI:** 10.1101/2022.06.24.497549

**Authors:** Jonathan M. Philpott, Alfred M. Freeberg, Jiyoung Park, Kwangjun Lee, Clarisse G. Ricci, Sabrina R. Hunt, Rajesh Narasimamurthy, David H. Segal, Rafael Robles, Yao D. Cai, Sarvind Tripathi, J. Andrew McCammon, David M. Virshup, Joanna C. Chiu, Choogon Lee, Carrie L. Partch

## Abstract

PERIOD (PER) and Casein Kinase 1δ regulate circadian rhythms through a phosphoswitch that controls PER stability and repressive activity in the molecular clock. CK1δ phosphorylation of the Familial Advanced Sleep Phase (FASP) serine cluster embedded within the Casein Kinase 1 binding domain (CK1BD) of mammalian PER1/2 inhibits its activity on phosphodegrons to stabilize PER and extend circadian period. Here, we show that the phosphorylated FASP region (pFASP) of PER2 directly interacts with and inhibits CK1δ. Co-crystal structures in conjunction with accelerated molecular dynamics simulations reveal how pFASP phosphoserines dock into conserved anion binding sites near the active site of CK1δ. Limiting phosphorylation of the FASP serine cluster reduces product inhibition, decreasing PER2 stability and shortens circadian period in human cells. We found that *Drosophila* PER also regulates CK1δ via feedback inhibition through the phosphorylated PER-Short domain, revealing a conserved mechanism by which PER phosphorylation near the CK1BD regulates CK1 kinase activity.

## Introduction

The mammalian circadian clock is driven by a set of interlocked transcription-translation feedback loops (TTFLs) (Takahashi, 2017). The core feedback loop is driven by the heterodimeric transcription factor, CLOCK:BMAL1, that promotes the transcription of multiple clock-controlled genes including its own repressors, Period (PER) and Cryptochrome (CRY) (Koike et al., 2012). PER proteins nucleate formation of a complex with CRYs and the clock-associated kinase, Casein Kinase 1 δ/ε (CK1) (Aryal et al., 2017; Lee et al., 2001; Michael et al., 2017) that ultimately bind to and inhibit CLOCK:BMAL1 activity (Cao et al., 2021), so the relative abundance of PER proteins is tightly regulated (Chen et al., 2009; Lee et al., 2011b) by transcriptional, post-transcriptional, and post-translational mechanisms (Crosby and Partch, 2020). The abundance of PER proteins is critical, as constitutive overexpression of PER proteins disrupts circadian rhythms (Chen et al., 2009) while the inducible regulation of PER expression on a daily basis can establish circadian rhythms *de novo* with tunable periods (D’Alessandro et al., 2015).

The post-translational control of PER stability by its cognate kinase, CK1, has been widely studied. CK1 remains stably bound, or anchored, to PER1 and PER2 throughout the circadian cycle (Aryal et al., 2017; Lee et al., 2001) via a conserved Casein Kinase 1 Binding Domain (CK1BD) (Eide et al., 2005; Lee et al., 2004). CK1 regulates PER2 stability through a phosphoswitch mechanism, whereby the anchored kinase phosphorylates antagonistic sites on PER2 that, on balance, control its stability (Zhou et al., 2015). CK1 phosphorylation of a Degron located several hundred residues upstream of the CK1BD, near the tandem PAS domains of PER2, leads to recruitment of the E3 ubiquitin ligase, β-TrCP, and subsequent proteasomal degradation (Eide et al., 2005; Masuda et al., 2020; Ohsaki et al., 2008; Vanselow et al., 2006). Activity at this PAS-Degron is counteracted somehow by CK1-dependent phosphorylation of a multiserine cluster within the CK1BD known as the FASP region. This region is named for a Ser to Gly polymorphism in human *Per2* (S662G) that disrupts CK1 activity at this cluster (Narasimamurthy et al., 2018), destabilizing PER2 and shortening circadian period, leading to Familial Advanced Sleep Phase Syndrome (Toh et al., 2001) that impacts the timing of sleep onset in humans (Jones et al., 2013). Mutations in CK1 can also alter the balance of its activity on these two regions to significantly shorten circadian period *in vivo* (Lowrey et al., 2000; Xu et al., 2005; Xu et al., 2007).

Similar to the FASP region of mammalian PER1/2, phosphorylation of the PER-Short domain of *Drosophila* PER (dPER) by DOUBLETIME (DBT, the *Drosophila* homolog of CK1δ/ε) and NEMO/NLK attenuates DBT-dependent phosphorylation at an N-terminal Degron to stabilize dPER and regulate circadian period (Chiu et al., 2011; Chiu et al., 2008). In particular, DBT-dependent phosphorylation of S589, the site of the classic *per*^S^ mutation (S589N) within the PER-Short domain (Konopka and Benzer, 1971), stabilizes PER in its inhibition-active form while mutation of this residue is sufficient to destabilize PER and shorten circadian period (Top et al., 2018). Phosphorylation of S596 by the NEMO/NLK kinase is required for DBT-dependent phosphorylation of nearby sites within the PER-Short domain, including S589, suggesting a hierarchical phosphorylation program that acts as a time-delay in the regulation of dPER abundance and repressive activity (Chiu et al., 2011).

In this study, we discovered how CK1 phosphorylation of the stabilizing FASP or PER-Short domains leads to feedback inhibition of the kinase. First, we found that multisite phosphorylation of the human PER2 FASP region is gated by rate-limiting phosphorylation of the first of five sequential serines in the FASP cluster by CK1. After this slow priming, CK1 rapidly phosphorylates the remaining serines in the FASP cluster following an ordered-distributive kinetic mechanism. Phosphorylation of C-terminal sites within the FASP region leads to product inhibition of CK1, reducing the overall rate of priming phosphorylation. Consistent with this, we show that phosphorylated FASP (pFASP) peptides inhibit CK1 activity on the PAS-Degron *in vitro.* Crystal structures and molecular dynamics simulations of the pFASP-bound kinase reveal the mechanism of product inhibition, driven by the interaction between pFASP and highly conserved anion binding sites on CK1. Leveraging newly developed U2OS *PER1^Luc^*and *PER2^Luc^* reporter cell lines (Park, 2022), we used CRISPR-generated indels to disrupt phosphorylation within the FASP region of *Per1* or *Per2*, observing shorter circadian periods and demonstrating conservation of the pFASP phosphoswitch in PER1 and PER2. Likewise, we discovered that phosphorylation of the PER-Short domain of *Drosophila* PER, located just upstream of the CK1BD, also inhibits CK1 activity *in vitro.* A structure of CK1 bound to a PER-Short domain phosphopeptide reveals a similar mechanism of product inhibition. Taken together, these results establish a conserved mechanism by which CK1 activity is regulated by feedback inhibition of PER to control PER stability and kinase activity within the molecular clock.

## Results

### CK1 phosphorylates the serine cluster in the PER2 FASP region in a sequential manner

The FASP region is an intrinsically disordered stretch of ∼50 highly conserved residues embedded within the CK1BD of PER1 and PER2 that contains a cluster of 5 serine residues with the repeated spacing of SxxS (**Figure 1a**). We previously showed that CK1 is necessary and sufficient to initiate phosphorylation of the FASP region in mouse PER2 by targeting the first serine within this cluster in a rate-limiting step known as priming (Narasimamurthy et al., 2018). We sought to extend this analysis to study the mechanism of multisite phosphorylation within the intact human PER2 FASP region using an NMR-based kinase assay that provides site-specific resolution of kinase activity on an ^15^N-labeled peptide substrate (Narasimamurthy et al., 2018; Smith et al., 2015). Using a constitutively active version of CK1δ lacking its autoinhibitory C-terminal tail, we found that CK1 phosphorylated all 5 serines within the FASP region (**Figure 1b**, **Supplementary Figure 1a**). Based on the CK1 consensus recognition motif, pSxxS (Flotow et al., 1990), we expected that CK1 would phosphorylate serines downstream of the priming site in a sequential manner. To test this, we introduced alanine mutations at each successive serine within the FASP cluster and assayed CK1 activity in the NMR kinase assay (**Figure 1c-f**). As expected, each Ser/Ala substitution selectively disrupted CK1 activity at downstream serines, confirming that the kinase acts in a sequential manner and generating a set of variably phosphorylated FASP peptides to study the functional consequences of kinase activity on this region.

**Figure 1.**
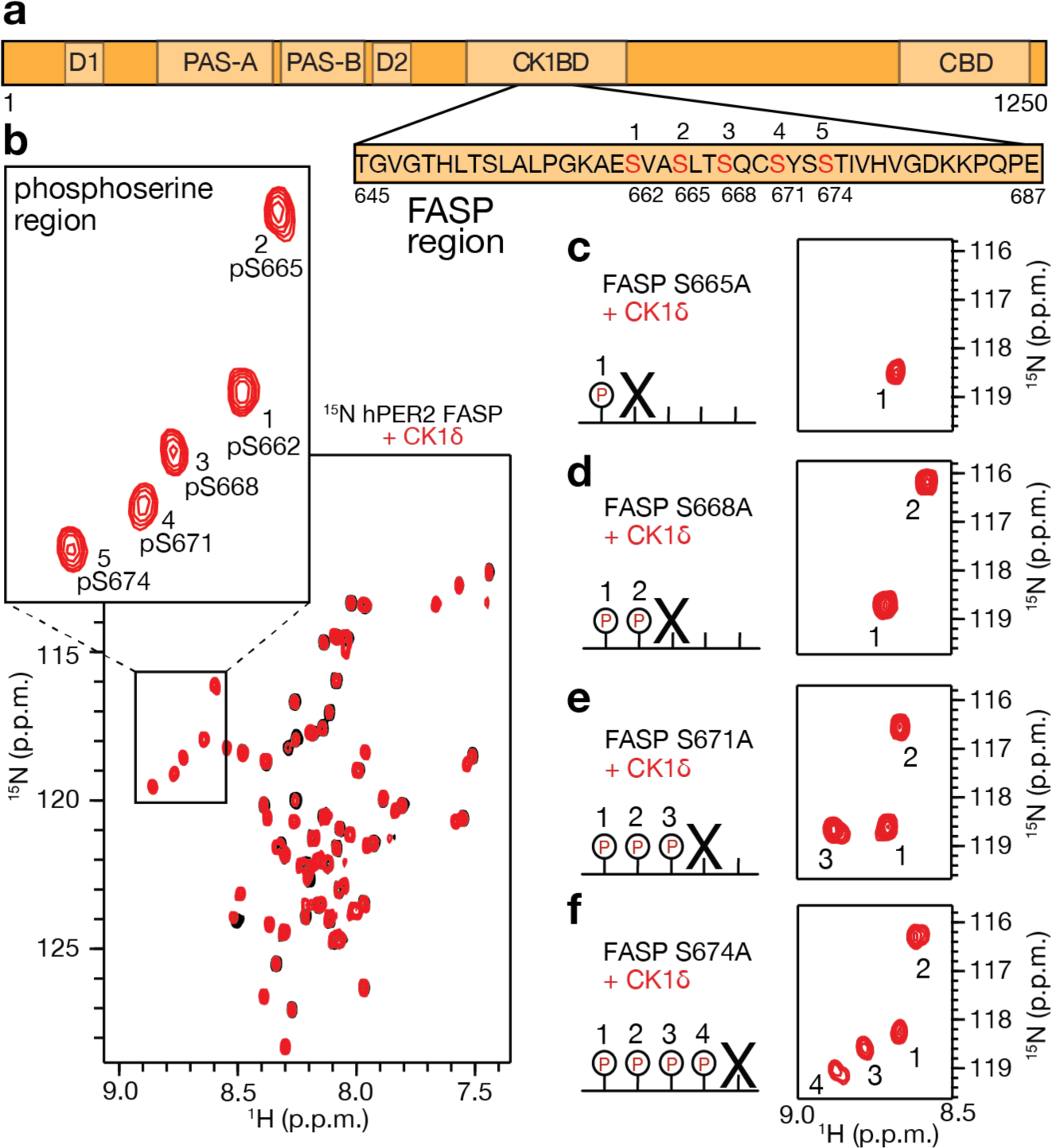
CK1 phosphorylates the human PER2 FASP region sequentially. **a**, Domain map of hPER2 with tandem PAS domains, phosphodegrons (D1/D2), Casein Kinase 1 binding domain (CKBD), and CRY binding domain (CBD). Zoom, the FASP peptide used in NMR assays with CK1 phosphorylation sites (red). **b**, ^15^N/^1^H HSQC of 200 µM hPER2 FASP used for kinetic assays (black), overlayed with 16 hr timepoint in kinase assay (red). Zoom/boxed region, the phosphoserine region. Numbering (1-5) corresponds to the order in the FASP peptide. **c-f**, Schematic representation of Ser/Ala mutations introduced to the FASP peptide. X, indicates position of the alanine substitution within the FASP serine cluster. Zoom of phosphoserine region for the ^15^N/^1^H HSQC of FASP mutants (red) at 3 hr timepoint in the kinase assay, with numbering showing the sequential order of reaction.

### CK1 follows an ordered distributive mechanism gated by slow, non-consensus priming

Due to the sequential nature of FASP phosphorylation by CK1, we propose that the kinase follows an ordered distributed kinetic mechanism (**Figure 2a**). The transient accumulation of intermediates can theoretically be observed by NMR, depending on the rate constants for intermediate states (**Figure 2b-c**) (Cordier et al., 2012). Since we previously determined that phosphorylation of the non-consensus priming serine was much slower than consensus-based activity on subsequent serines (Narasimamurthy et al., 2018), we expected that phosphorylation of the FASP would appear to be an all-or-none event by NMR, represented by state F in **Figure 2b**. When phosphorylation sites are in close proximity to one another, the chemical shift of a particular residue can change over the course of a reaction as nearby sites become modified (Smith et al., 2015). Using the distinct phosphospecies generated with alanine substitutions in the FASP, we were able to assign unique chemical shifts for each phosphopeak that arises as a function of sequential phosphorylation (**Supplementary Figure 1b**). However, in a kinase reaction with the native FASP, we did not observe accumulation of any intermediate states over the course of the reaction, only a set of peaks corresponding to the 5 phosphoserines defined as species C in **Figure 2d** and **e**. We did observe a second slow step due to phosphorylation of T675 (peak 6D in **Figure 2d**) that was dependent on phosphorylation of the upstream residue S671 (**Supplementary Figure 1b**), consistent with the less optimal CK1 consensus motif of pSxxxS/T (Flotow et al., 1990). The peaks corresponding to phosphoserines 4 and 5 (pS671 and pS674) were split depending on whether or not T675 was phosphorylated (4C and 4D, 5C and 5D, **Figure 2d**).

**Figure 2.**
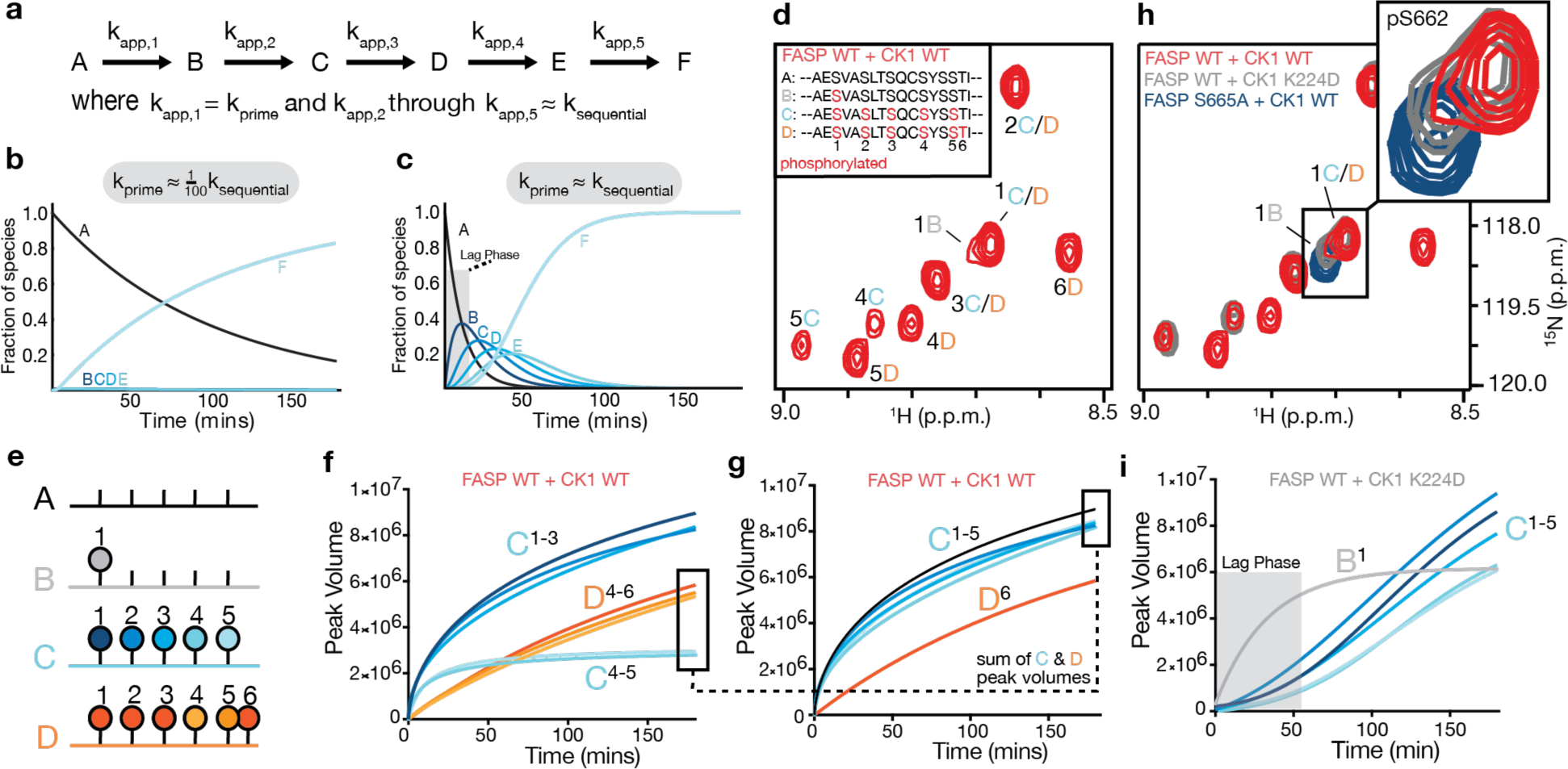
CK1 follows an ordered distributive mechanism on the human PER2 FASP region. **a**, Schematic of ordered distributive kinetic model used for panels **b** and **c** (modeled using Wolfram Mathematica, see Supplement). **b**, Reaction coordinate for ordered distributive kinetic mechanism with differential rates for priming and sequential kinase activity. **c**, Reaction coordinate for ordered distributive kinetic mechanism with similar rates for priming and sequential kinase activity. **d**, Zoom of ^15^N/^1^H HSQC hPER2 FASP phosphoserine region looking at peaks corresponding to different states arising from at 3 hr incubation with CK1 WT (red). States A-D (inset) correspond to unique chemical shift states observed throughout the kinase assay. **e**, Schematic of unique states A-D observed in **d**: A, unphosphorylated FASP; B, primed FASP; C, all serines in FASP phosphorylated; D, all serines plus T675 phosphorylated. Numbering indicates order of phosphorylation. **f**, Traces of accumulating peak volume from the NMR kinase assay. Letters C-D with superscript numbers correspond to specific phosphoserines observed in the NMR assay. **g**, Same traces as in **f**, with peak volumes corresponding to states C and D for the last two serines (4-5) summed to show that all phosphoserines in the FASP report on state C with similar kinetics. **h**, Phosphoserine region showing unique chemical shift environment for singly phosphorylated (priming only) FASP S665A (blue) overlayed with WT FASP with CK1 WT (red) or K224D (gray) after 3 hr incubation. B-D lettering corresponds to unique states observed in NMR kinase assay as in **d**. **i**, Traces of accumulating peak volume from NMR kinase assay with CK1 K224D, resolving the transient accumulation of primed FASP and a clear lagging phase for subsequent phosphorylation states.

We confirmed the overall kinetic scheme by collecting a series of 6-minute SoFast HMQC NMR spectra over the course of a 3-hr real-time kinase reaction in the magnet (**Figure 2f**). Combining the total intensities for phosphoserines 4 and 5 from states C and D (**Figure 2g**) confirms that the reaction kinetics proceed as predicted by the model outlined in **Figure 2b**. To probe the ordered distributive mechanism, we utilized a mutant form of the kinase, K224D (Shinohara et al., 2017), that disrupts the anion binding pocket Site 1 near the active site thought to anchor primed substrates and facilitate kinase activity on downstream consensus sites (Longenecker et al., 1996; Venkatesan et al., 2019; Zeringo and Bellizzi, 2014). The K224D mutant retains its ability to prime the FASP region with kinetics similar to the WT kinase but has decreased activity on subsequent consensus-based sites (Philpott et al., 2020). By reducing the relative difference in rates for priming and sequential phosphorylation with this mutant, we were able to resolve a distinct peak corresponding to the transient accumulation of the primed, singly phosphorylated FASP, like that observed in the PER2 S665A mutant (**Figure 2h**), which exhibited a clear lagging phase for kinase activity on subsequent phosphorylation sites (**Figure 2i**).

Mutation of the priming serine in the human FASP region to an aspartate, S662D, rescues downstream kinase activity within the FASP region (Toh et al., 2001) and increases circadian period by promoting the stabilization of PER2 in mice (Xu et al., 2007). Other CK1 substrates are constitutively primed by D/E residues upstream that can act as phosphomimetics (Flotow and Roach, 1991; Marin et al., 2003). Using the NMR kinase assay, we observed that while the S662D mutation can prime kinase activity on downstream serines within the FASP region, it is a relatively poor mimetic of pS662, leading to a reduction in the overall kinase activity on the FASP peptide (**Supplementary Figure 1c-d**). Therefore, the ability of the S662D mutant to promote PER2 stability and period lengthening *in vivo* (Xu et al., 2007) likely rests on the weak but constitutive priming by the aspartate.

### Sequential phosphorylation of FASP leads to feedback inhibition of CK1

Disrupting phosphorylation of the FASP region by mutating the priming serine increases CK1 activity at a phosphodegron in PER2 (Philpott et al., 2020), consistent with increased PER2 turnover in the human FASPS S662G mutant (Toh et al., 2001). However, no mechanistic model has been robustly demonstrated yet for how FASP phosphorylation influences CK1 activity. Monitoring both the loss of peak intensity for S662 and the concomitant increase in peak intensity for pS662, we made the striking observation that the fraction of primed WT FASP plateaued at approximately 50% completion over the course of a 3-hr real-time NMR kinase assay (**Figure 3a**). This suggested to us that some or all of the phosphoserines in the FASP cluster might work to reduce CK1 activity via feedback inhibition. To dissect the role of successive phosphoserines in this inhibition, we monitored the kinetics of FASP priming by NMR using the alanine-substituted mutants that limit the extent of sequential phosphorylation in a stepwise manner (**Figure 3b-e**). We found that reducing the extent of possible phosphosites in the FASP led to a stepwise increase in priming activity, resulting in near complete priming of the S668A mutant limited to just 2 phosphoserines (**Figure 3d**) and a several-fold increase in the rate of priming (k_prime_) relative to the WT FASP (**Figure 3f**).

**Figure 3.**
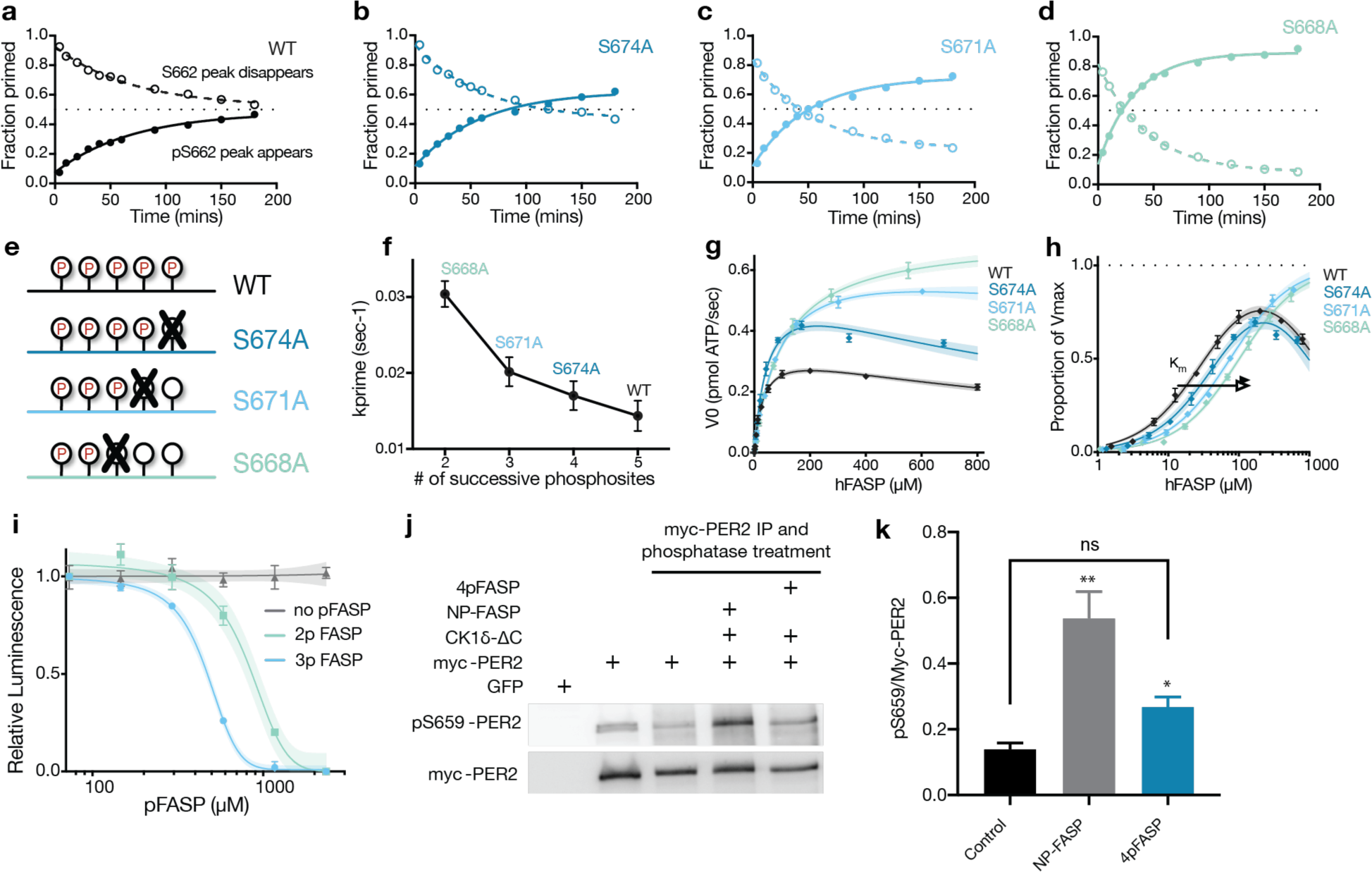
Phosphorylation of the human PER2 FASP region inhibits CK1 activity. **a**-**d**, NMR kinase assay for the WT FASP or indicated mutant peptides monitoring the reaction kinetics of priming phosphorylation at S662 by NMR. **e**, Schematic of FASP alanine mutations and resulting discrete phosphostates. **f**, Plot of priming rate constant (k_prime_) as a function of the possible successive phosphosites in the FASP. Error bars represent SEM from fits in panels **a-d**. **g**, ADP-Glo kinase assay with titration of FASP mutant peptides with mean and SD from 2 replicates, representative of n = 3 independent assays. Shaded area indicates 95% C.I. of the fit. **h**, Data from panel **f** normalized by V_max_ values calculated from the preferred kinetic model (see Supplementary Figure 2, Table 1). **i**, ADP-Glo kinase assay of hPER2 PAS-degron peptide (see Supplementary Figure 2b) with titration of pFASP peptides corresponding to 2 (2p) or 3 (3p) phosphoserines (see Supplementary Figure 4c) with mean and SD from 2 replicates, representative of n = 3 independent assays. Shaded area indicates 95% C.I. of the fit. **j-k**, Western blot and quantification of the phosphorylation of the FASP priming site (pS569 in mouse PER2). Full-length mouse PER2 was immunoprecipitated from transfected HEK293T cells, dephosphorylated, and subjected to an *in vitro* kinase assay with 200 ng CK1 in the presence and absence of added NP or 4pFASP peptides as indicated.

**Table 1.** X-ray crystallography data collection and refinement statistics.

|  | 2pFASP:CK1δ | 3pFASP:CK1δ | 4pFASP:CK1δ | pS589:CK1δ |
| --- | --- | --- | --- | --- |
| **Data collection** |  |  |  |  |
| PDB Id. | 8D7M | 8D7N | 8D7O | 8D7P |
| Beam line | APS(23IDD) | APS(23IDD) | APS(23IDD) | APS(23IDD) |
| Resolution Range (Å)<br>(highest shell)* | 46.15 - 2.25<br>(2.32 - 2.25) | 68.01-1.66<br>(1.69 - 1.66) | 67.70 - 1.65<br>(1.68 - 1.65) | 60.77 – 2.25<br>(2.32 - 2.25) |
| Space group | C 1 2 1 | C 1 2 1 | C 1 2 1 | P1 |
| a, b, c | 55.7, 135.97,<br>91.14 | 55.90, 136.03,<br>90.54 | 55.33, 135.40,<br>90.35 | 48.78, 56.74,<br>65.94 |
| $\alpha, \beta, \gamma$ | 90, 94.3, 90 | 90, 94.60, 90 | 90, 94.41, 90 | 108.93, 95.63,<br>108.69 |
| Wavelength (Å) | 1.03 | 1.03 | 1.03 | 1.03 |
| Total observations | 176957 (15319) | 480416 (23861) | 468136 (24113) | 85303 (7506) |
| Unique reflections | 31319 (2767) | 79274 (3942) | 78946 (4059) | 27457 (2505) |
| Completeness (%) | 97.8 (94.5) | 100 (99.9) | 99.4 (99.2) | 93.7 (91.8) |
| R<sub>merge</sub> | 15.2 (127.4) | 9.3 (55.6) | 6.4 (54.2) | 11.6 (31.2) |
| <I/σ> | 9.4 (2.5) | 9.9 (2.7) | 13.6 (2.9) | 5.5 (2.7) |
| CC1/2 | 0.99 (0.55) | 0.99 (0.82) | 0.99 (0.84) | 0.98 (0.85) |
| Redundancy | 5.7 (5.5) | 6.1 (6.1) | 5.9 (5.9) | 3.1 (3.0) |
| **Refinement** |  |  |  |  |
| R<sub>work</sub>/ R<sub>free</sub> (%) | 17.9 (22.9) | 18.8 (21.3) | 17.4 (21.1) | 19.7 (25.4) |
| Number of non-hydrogen atoms | 5075 | 5388 | 5342 | 4999 |
| Protein | 4820 | 4840 | 4840 | 4864 |
| Water | 255 | 548 | 502 | 135 |
| B-factor (Wilson) | 36.95 | 33.32 | 32.9 | 29.93 |
| RMSD Bond length (Å) | 0.008 | 0.01 | 0.011 | 0.009 |
| RMSD Bond angle | 1.05 | 1.18 | 1.18 | 1.04 |
| Ramachandran favored (%)/<br>Ramachandran outliers (%) | 97.04/0.0 | 96.17/0.0 | 96.34/0.26 | 96.23/0.0 |

To confirm that the observed inhibition of FASP priming was due to feedback inhibition, we performed substrate titration experiments using an ADP-Glo assay that measures bulk kinase activity. Here, we observed that WT FASP had the lowest overall rate of phosphorylation, despite having the most consensus-based phosphosites available for the kinase (**Figure 3e,g**). Moreover, kinase activity began to decrease on WT FASP at higher substrate concentrations, consistent with substrate/product inhibition. As in the NMR based assay, we observed a rank-order increase in overall kinase activity and decrease in apparent inhibition as the extent of possible phosphosites in FASP was successively limited with the alanine substitutions (**Figure 3g**), shifting the Michaelis constant higher for each of these peptides (**Figure 3h**, **Supplementary Figure 2a**). Together, these data are consistent with the rapid generation of a pFASP product that inhibits kinase activity in *trans*. To test this, we performed a kinase assay on a peptide corresponding to the PAS-Degron of PER2 (**Supplementary Figure 2b**) (Isojima et al., 2009) in the presence or absence of synthetically phosphorylated FASP peptides (pFASP). The doubly phosphorylated FASP peptide (2pFASP) inhibited the kinase in *trans*, with the 3pFASP peptide further increasing the potency of inhibition (**Figure 3i**). Addition of the 4pFASP peptide in *trans* also inhibited phosphorylation of the priming serine in an *in vitro* kinase assay, where full-length mouse PER2 immunoprecipitated from HEK293 cells was dephosphorylated *in vitro* and then treated with CK1δ kinase that had been pre-incubated with 4pFASP peptide (**Figure 3j-k**).

We performed real-time bioluminescence measurements of HEK293T cells transiently transfected with hPER2::LUC and CK1δ expression plasmids to monitor the half-life of hPER2::LUC in the context of Ser/Ala mutations that modulate PER2 phosphorylation. Consistent with prior results, mutation of the PAS-Degron site (S480) to an alanine increased hPER2::LUC stability (**Supplementary Figure 2c**), where S480 corresponds to the first phosphorylated residue of the β-TrCP consensus recognition motif, DpSGYGpS (**Supplementary Figure 2b**) (Eide et al., 2005; Reischl et al., 2007; Wu et al., 2003). Mutation of either the priming serine (S662A) or downstream serines (i.e., S671A or S674A) decreased PER2::LUC half-life, while mutation of the threonine residue at the end of the FASP cluster (T675A) showed no significant effect on PER2 stability (**Supplementary Figure 2c**). To further link the Ser/Ala mutations in FASP to β-TrCP recognition of PER2, we performed co-immunoprecipitation experiments from HEK293T cells co-transfected with PER2, β-TrCP and CK1δ expression plasmids and observed increased CK1δ-dependent β-TrCP binding to PER2 with mutants that reduce FASP phosphorylation (**Supplementary Figure 2d**).

### pFASP binds to CK1 via conserved anion binding sites to occlude the substrate binding cleft

CK1 has several highly conserved anion binding sites located around the substrate binding cleft (**Figure 4a**) (Longenecker et al., 1996; Philpott et al., 2020). Given their location and the distance between phosphosites within pFASP, we hypothesized that these anion binding sites might mediate sinteraction of the phosphorylated peptide with CK1. To test this prediction, we solved crystal structures of phosphorylated FASP peptides bound to the catalytic domain of CK1δ (**Table 1**). Indeed, in three distinct complexes with 2pFASP, 3pFASP, or 4pFASP peptides, we observed a consistent binding mode (**Supplementary Figure 3a-b**) with the phosphorylated peptides coordinating the two anion binding sites (Site 1 and Site 2) to fully occlude the substrate binding cleft (**Figure 4b**). Interestingly, a substrate motif analysis derived from a dataset of 101 known CK1δ substrate sequences from PhosphoSitePlus aligns well with this pFASP binding mode, suggesting that the first 3 sites of the FASP region conform to an ideal CK1δ recognition sequence (**Supplementary Figure 3c**). In Site 1, R178 and the backbone amide of G215 coordinate pS662 of the pFASP peptide (**Figure 4c**). Site 1 harbors the location of the *tau* mutation (R178C) that shortens circadian period (Lowrey et al., 2000). We did not observe strong density for K224, the other basic residue that could coordinate an anion in Site 1, suggesting flexibility in the Fɑ helix consistent with previous studies (Cullati et al., 2022; Philpott et al., 2020; Shinohara et al., 2017). Moving down the pFASP peptide, we observed additional backbone-backbone interactions and docking of the pFASP side chains of V663 & A664 into small hydrophobic pockets within the substrate binding cleft (**Figure 4b, d**). We did not observe strong density for K224, the other basic residue that could coordinate an anion in Site 1, suggesting flexibility in the Fɑ helix consistent with previous studies (Cullati et al., 2022; Philpott et al., 2020; Shinohara et al., 2017). Moving down the pFASP peptide, we observed additional backbone-backbone interactions and docking of the pFASP side chains of V663 & A664 into small hydrophobic pockets within the substrate binding cleft (**Figure 4b, d**).

**Figure 4.**
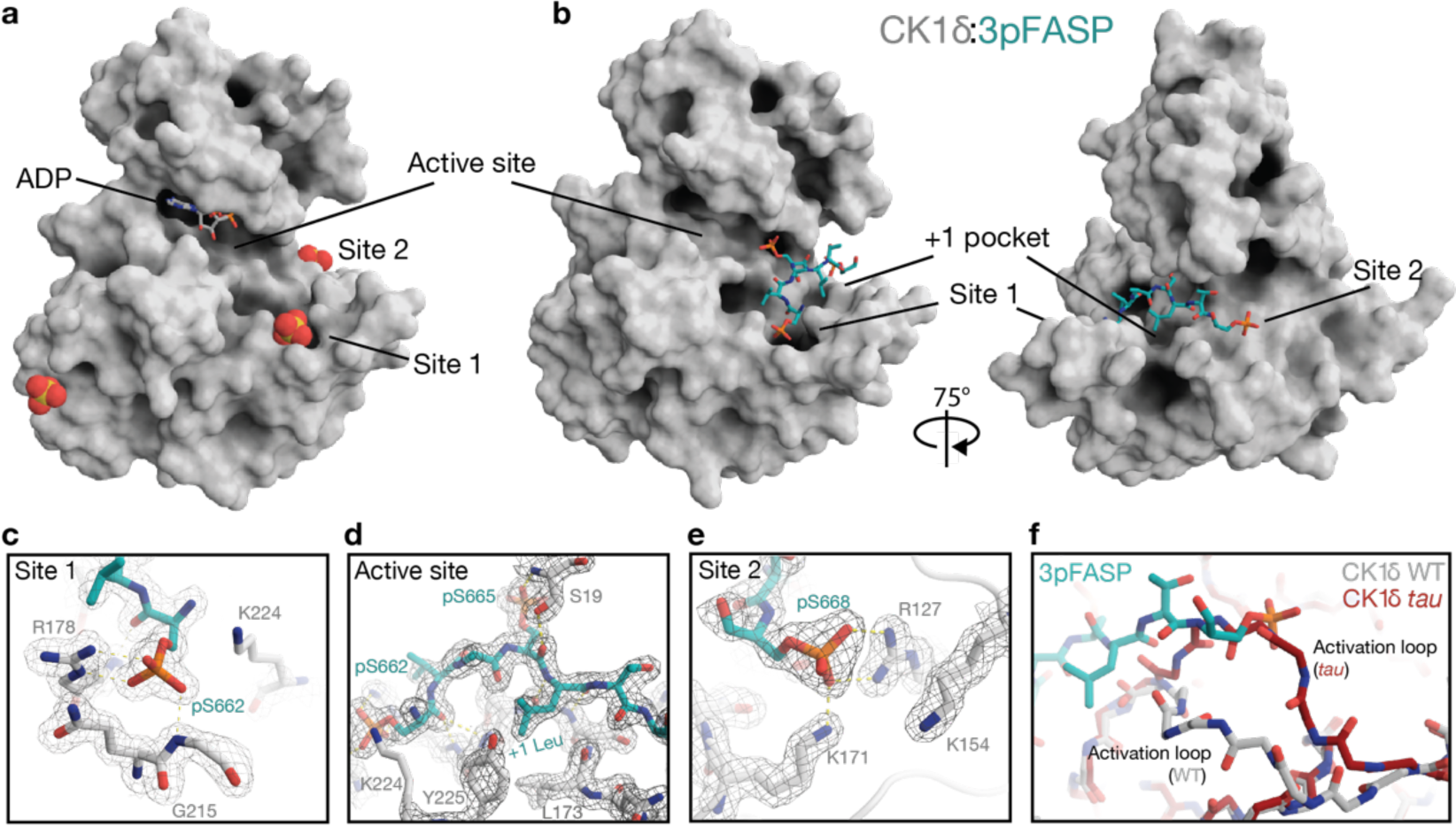
The human PER2 pFASP binds to the active site of CK1. **a**, Surface representation of CK1 catalytic domain bound to ADP (PDB 5X17). Spheres, sulfate anions from the crystallization condition bound at anion binding sites as indicated. **b**, Surface representation of the CK1 catalytic domain bound to a 3pFASP peptide (see Supplementary Figure 4). **c**, Zoom of 3pFASP interactions within anion binding site 1. **d**, Zoom of 3pFASP interactions within active site region. **e**, Zoom of 3pFASP interactions within anion binding site 2. **f**, Structural alignment showing the main chain for the activation loop of CK1 WT (gray) and the *tau* mutant (red) showing a clash of the 3pFASP (teal) with the conformation of the activation loop stabilized by the *tau* mutation.

The 2^nd^ phosphoserine of pFASP, pS665, projected into the active site of CK1, positioned close to where the gamma phosphate would be in ATP-bound CK1 (**Supplementary Figure 3e**). We also observed that S19 of the N-terminal lobe P-loop of CK1 clamped down on pS665, suggesting a stabilizing mechanism for transfer of phosphate from the nucleotide to the substrate, with the +1 Leu after pS665 fitting into a small hydrophobic pocket on the kinase. As previously noted, this small hydrophobic pocket is formed between L173 and Y225 of CK1 (the+1 pocket, **Figure 4b, d**) in the downward conformation of the CK1 activation loop (Philpott et al., 2020). The +1 pocket is just large enough to accommodate small residues such as Ala, Leu, or Val, and could contribute to CK1 recognition of the non-consensus SLS motif (Marin et al., 2003) or the FASP priming site, where the +1 residue is a valine.

Moving towards the C-terminal end of the peptide, the 3^rd^ phosphoserine of pFASP, pS668, is coordinated by anion binding Site 2, comprising basic residues R127, K154, and K171 (**Figure 4e**). R127 is part of the conserved HRD motif involved in the regulation of Ser/Thr kinases by coordinating a phosphorylated residue within the activation loop (Nolen et al., 2004). To further validate the significance of these anion binding sites in the binding of pFASP and product inhibition, we introduced charge inversion mutations in Site 1 (K224D) or Site 2 (R127E) and performed substrate titrations with FASP peptide, observing a decrease in the level of product inhibition (**Supplementary Figure 3d**).

The highly specific sequential mechanism of FASP phosphorylation also strongly suggests that the FASP peptide translates through the substrate binding cleft, with each phosphoserine anchoring into Site 1 to facilitate the next phosphorylation event in the active site. The phosphorylated serines in pFASP all share the same approximate atomic distance between each other that is consistent with the spacing between Site 1, the active site, and Site 2. Therefore, product inhibition may continue to increase as a function of sequential phosphorylation at least partially due to changes in entropy, where 3pFASP can occupy each of the aforementioned sites on the kinase in one way (1 microstate), 4pFASP in two ways (2 microstates), and 5pFASP in three ways (3 microstates) (**Supplementary Figure 3f**).

The *tau* mutation in Site 1 of CK1δ prevents anion coordination at this site and alters the global dynamics of CK1 as well as significantly changing the local dynamics in and around the activation loop and Site 2 via an allosteric mechanism (Philpott et al., 2020). We note that the alternate conformation of the activation loop stabilized in the *tau* mutant sterically clashes with the 3pFASP binding mode (**Figure 4f**). This incompatibility suggests that the *tau* mutant is less susceptible to feedback product inhibition, perhaps also contributing to the increased phosphorylation of the PAS-Degron and degradation of PER2 by this kinase mutant (Gallego et al., 2006a; Philpott et al., 2020; Zhou et al., 2015).

### Molecular dynamics simulations support stable binding mode of pFASP:CK1 crystal structures

To investigate how CK1 might be inhibited by a fully phosphorylated FASP peptide, we modeled and simulated the interaction of CK1 and a 5pFASP peptide using Gaussian accelerated Molecular Dynamics (details in Supplementary Material) (Miao et al., 2015). We found that the first three phosphoserines (pS662, pS665, and pS668) remain stably bound in the substrate binding cleft of CK1, while the 4^th^ and 5^th^ phosphoserines (pS671 and pS674) display higher mobility (**Figure 5a-b**). As expected, pS662 forms stable electrostatic interactions in Site 1 (**Figure 5c**). In addition to being close to R178 and K224, pS662 also engages in stable electrostatic interactions with Q214, located in the highly flexible loop-EF (Cullati et al., 2022; Shinohara et al., 2017). In the active site, pS665 is locked in place by a cluster of electrostatic interactions involving D128, K130, D148 and a Na^+^ ion (**Figure 5d**). The presence of this ion in the active site suggests that at least one cation (likely Mg^2+^) could be important for product inhibition. The third phosphoserine (pS668) is held in place in Site 2 by stable electrostatic interactions with R127, K154, K171, and, to a lesser extent, R168 (**Figure 5e**). Finally, we found that the 5^th^ phosphoserine (pS674) often engages in electrostatic interactions with R160 (close to anion binding Site 3, which precedes the activation loop) and with K155 (just after the DFG motif) (**Figure 5l**). We refer to this herein as Site 3’. These interactions provide additional product stabilization, perhaps contributing to stronger inhibition when the FASP region is fully phosphorylated.

**Figure 5.**
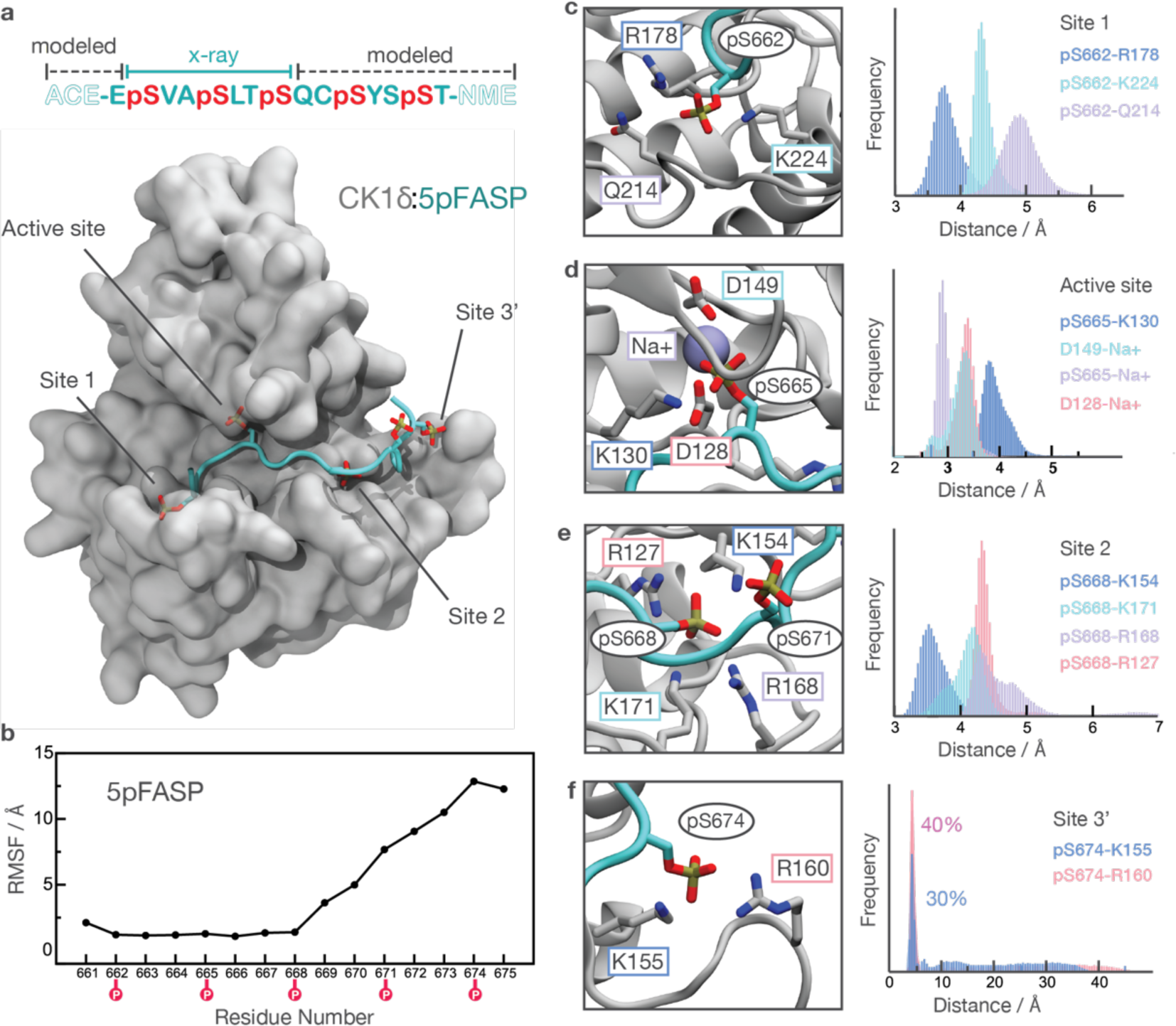
Molecular dynamics simulations of CK1:5pFASP. **a**, Structural model of CK1 bound to a 5p FASP peptide obtained by Gaussian accelerated Molecular Dynamics (GaMD). The molecular representation corresponds to one snapshot from the MD trajectories. **b**, Root Mean Square Fluctuation (RMSF) of the 5pFASP peptide with respect to its average conformation after aligning the MD trajectories with respect to CK1 backbone. **c-f**, Characterization of key interactions that stabilize the 5pFASP product in (**c**) Site 1, (**d**) the active site, (**e**) Site 2, and (**f**) an additional anion binding site located proximal to Site 2 (Site 3’). Histograms were computed based on distances sampled during the GaMD simulations.

### Circadian rhythms are shortened by small deletions in the conserved FASP region of PER

To address the regulatory role of the FASP region on circadian period, small amino acid deletions were introduced in the human *Per2* FASP region encoded by exon 17 (E17) by targeting the priming serine (S662) with CRISPR (**Figure 6a**). This CRISPR-mediated strategy utilized human U2OS cells with endogenous *Per2-luc* and *Per1-luc* knock-in reporter genes that produced robust rhythms in bioluminescence and clock proteins (Park, 2022). As out-of-frame mutations would disrupt LUC expression and eliminate bioluminescence, we could select for small in-frame deletions in PER2::LUC protein based on intact bioluminescence signals followed by molecular characterization. Small in-frame deletions that removed the priming serine led to rhythms in PER2::LUC abundance that were phase-advanced relative to WT PER2::LUC after synchronization by serum shock (**Figure 6b**). Additionally, PER1 also showed phase-advanced rhythms in abundance in these mutant lines relative to the parental cell line (**Supplementary Figure 4a**).

**Figure 6.**
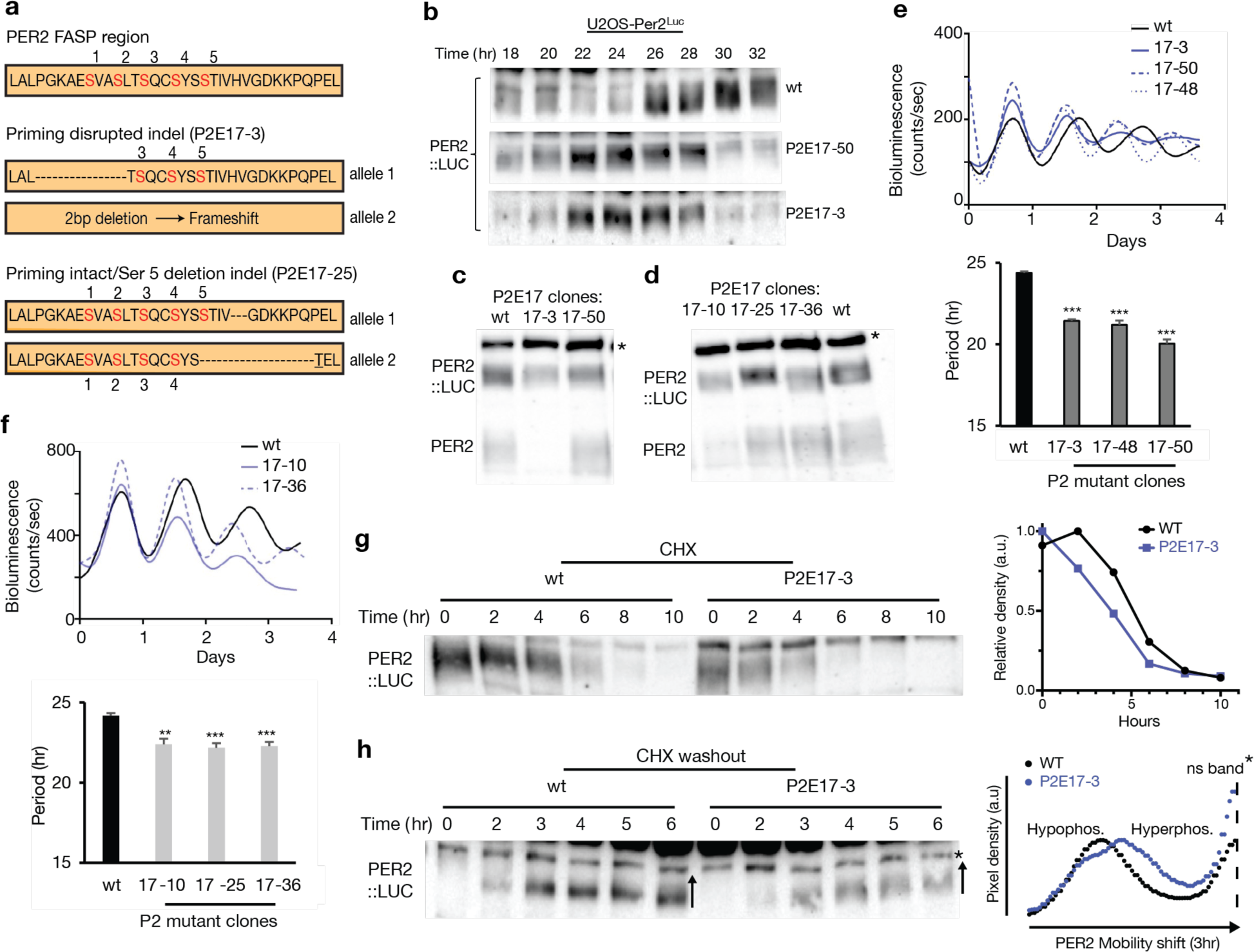
Circadian rhythms are shortened by small deletions in the conserved FASP domain of human PER2. **a**, Schematic representation of select in-frame deletions within the human PER2 FASP region are separated into 2 classes: priming-disrupted (P2E17-3) or priming intact (P2E17-25). **b**, Immunoblot of select priming-disrupted PER2::LUC mutants compared to WT PER2::LUC. Blot representative of two independent experiments. **c**,**d**, Representative immunoblots for (**c**) priming-disrupted or (**d**) priming intact PER2 mutants. *nonspecific band. The *Per2* allele in clone 17-3 has a frame-shifting mutation leading to deletion of the untagged PER2. **e**,**f**, Real-time bioluminescence traces of circadian rhythms from WT and mutant *Per2* clones (**e**, priming-disrupted; **f**, priming intact) with quantification of mean period and SD from n = 3 cultures. Periods from mutant clones were compared to WT with an unpaired t-test: ***, p < 0.001. *Per2* priming intact mutants (clones 17-25 and 17-36) exhibited rhythms that were shortened to a lesser degree than the priming-disrupted mutants in **e**, p < 0.05. In both graphs, the first peaks are aligned to show differences in period clearly. **g**, Western blot and quantification of degradation of WT PER2::LUC and clone 17-3 after cycloheximide treatment. Blot representative of two independent assays (n = 2). **h**, Western blot and quantification of phosphorylation of *de novo* PER2 after protein depletion by 10 hr CHX treatment and washout. Blot representative of two independent assays (n = 2).

Immunoblots for priming-disrupted PER2 mutants showed similar levels of gross phosphorylation as WT PER2::LUC (**Figure 6c-d**, **Supplemental Figure 4b**). To quantify differences in period, real-time bioluminescence measurements were collected on the clonal cell lines, which exhibited robust circadian rhythms with a significantly shorter period relative to the parental cell line (**Figure 6e**, **Supplemental Figure 4c**). Mutants that left the priming serine intact but removed one or more of the downstream serines (i.e., P2E17-25) also showed robust rhythms in bioluminescence with short periods (**Figure 6f**), but they were not as short as the priming-disrupted mutants that eliminate all FASP phosphorylation (P < 0.05). This suggests a role for the downstream phosphorylation sites in regulation of CK1 activity on PER2. Disruption of the FASP region led to decreased stability of PER2 after treatment with cycloheximide (**Figure 6g**) and also exhibited accelerated phosphorylation kinetics relative to WT, based on mobility shift, on the *de novo* PER2 synthesized after washout of cycloheximide (**Figure 6h**). The FASP region is highly conserved in PER1, and similar CRISPR-mediated edits to the PER1 FASP region by targeting the priming serine S714 also exhibited short period rhythms characterized by destabilized PER1 (**Supplemental Figure 5a-f**). NMR studies of the human PER1 FASP demonstrate that CK1 phosphorylates it similarly to PER2 and that indels disrupting the priming serine eliminate kinase activity on the region (**Supplemental Figure 5g**), demonstrating that the phosphorylated FASP region of both mammalian PERs likely act in a similar manner to constrain CK1 activity.

### The phosphorylated *Drosophila* PER-Short region binds CK1 to inhibit kinase activity

Mammalian and *Drosophila* PER proteins both possess the two conserved motifs comprising the CK1 binding domain (CK1BD) that stably anchors the kinase to PER throughout its daily life cycle (Eide et al., 2005; Kim et al., 2007; Kivimae et al., 2008; Nawathean et al., 2007) (**Figure 7a-b**). Phosphorylation of S589 on dPER by the *Drosophila* CK1 homolog DBT reduces kinase activity at a phosphodegron upstream to increase dPER stability and lengthen circadian period (Chiu et al., 2011; Kivimae et al., 2008) in a manner similar to the FASP-associated S662 on human PER2 (Toh et al., 2001) (**Figure 7a**). Loss of this phosphorylation site with the classic *per-Short* mutation (Konopka and Benzer, 1971) (*per*^S^, S589N) or other mutants nearby in the PER-Short domain result in predominantly short period phenotypes (Baylies et al., 1992; Chiu et al., 2011; Rothenfluh et al., 2000; Rutila et al., 1992). Because this CK1-dependent phosphosite is in close proximity to the CK1BD (**Figure 7c**) (Kim et al., 2007; Lee et al., 2004; Nawathean et al., 2007), we wondered if it would work similarly to the FASP region to attenuate CK1 activity. Phosphorylation of S589 by CK1 (i.e., DBT) has been invoked as a potential mechanism by which dPER could control CK1 activity in *trans* to regulate dPER turnover and repressive activity (Chiu et al., 2011; Chiu et al., 2008; Kivimae et al., 2008; Top et al., 2018). Phosphorylation of S589 is preceded *in vivo* by phosphorylation of S596 by NEMO kinase, and the S596A mutation eliminates phosphorylation of S589, suggesting a hierarchical regulation of the PER-Short region by multiple kinases (Chiu et al., 2011).

**Figure 7.**
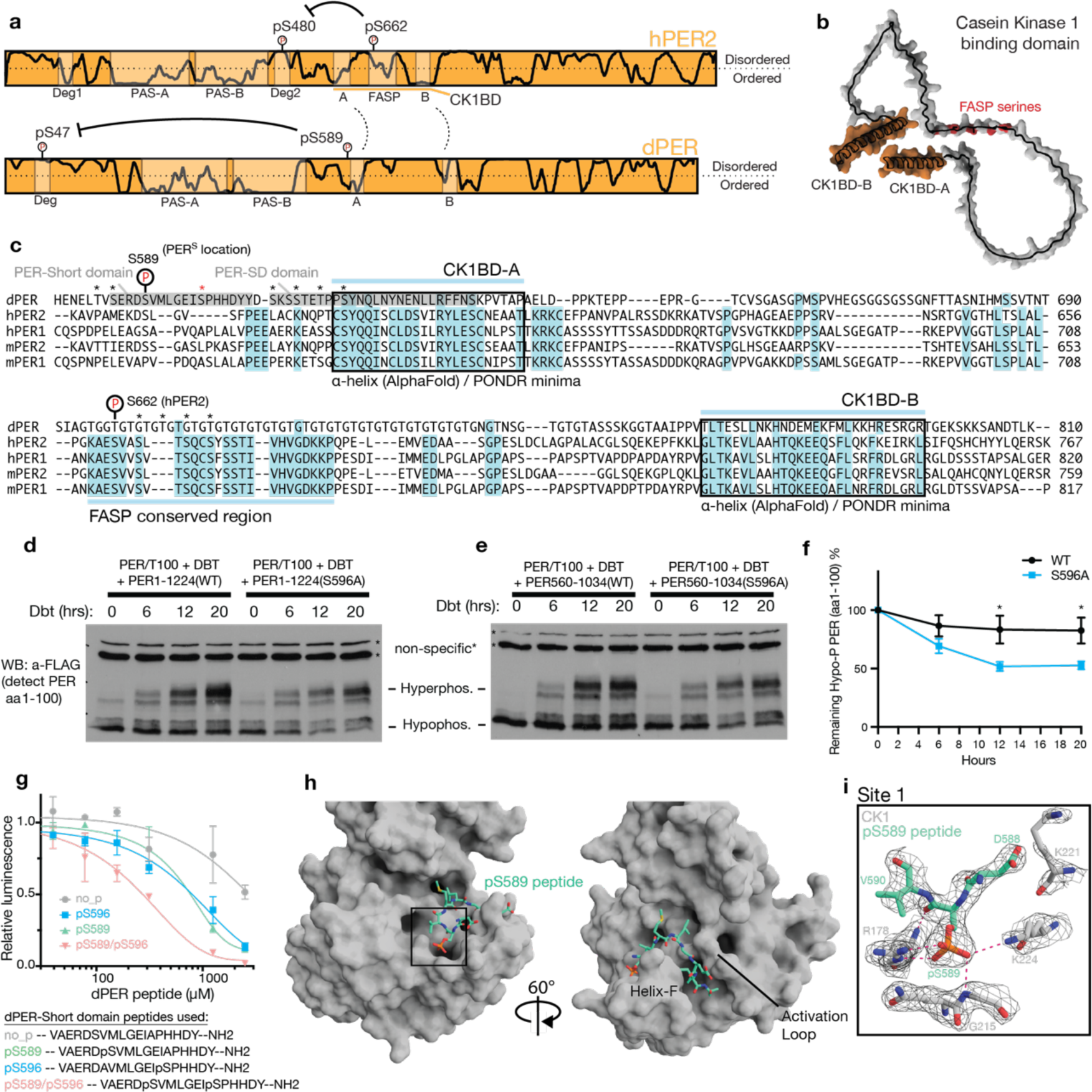
The phosphorylated PER-Short domain of *Drosophila* PER binds CK1 Site 1 to inhibit kinase activity. **a**, Schematic of human and *Drosophila* PER proteins, illustrating similar mechanisms of Degron regulation by phosphoserines proximal to the two conserved motifs of the CK1BD (see panel **c**). **b**, AlphaFold2 structural prediction of the human PER2 CK1BD with helical motifs A and B (orange) and FASP phosphorylation sites (red). **c**, Alignment of *Drosophila* PER, human and mouse PER1/2 proteins showing conservation and the dPER-Short and Short-Downstream (SD) domains. *, phosphorylation by either CK1 (black) or NEMO (red). **d-e**, Western blot of dPER fragment (aa1-100). Samples collected at indicated timepoints after kinase induction, followed by protein extraction and TEV cleavage. **f**, Quantification of hypophosphorylated band (n=4). WT and S596A compared with Two-way ANOVA with Sidak multiple comparisons test: *, p < 0.05. **g**, ADP-Glo kinase assay of CK1 on the human PER2 PAS-Degron substrate in the presence of indicated peptides from dPER. Data are mean and SD from 2 replicates, representative of n = 3 independent assays. **h**, Structure of human CK1 (gray) bound to the dPER-Short peptide with pS589 (light green). **i**, Close-up view of dPER pS589 coordinated by CK1 residues in anion binding Site 1.

To determine if the phosphocluster in the PER-Short domain regulates CK1 activity on the N-terminal dPER Degron, we examined the progressive phosphorylation of this region in *Drosophila* S2 cells using a full-length dPER construct that contains a TEV-cleavable 100-residue N-terminal fragment encompassing the Degron (PER/T100) (Chiu et al., 2011; Chiu et al., 2008). PER/T100 was co-expressed with full length dPER or a C-terminal fragment containing the kinase binding domain (residues 560-1034), with or without the S596A mutant, along with an inducible DBT construct. We observed that inhibition of dPER Degron phosphorylation is lost with the S596A mutant in full-length dPER (**Figure 7d, f**) or a fragment of dPER containing just the kinase binding domain in *trans* (**Figure 7e, f**). To test for direct inhibition of CK1 activity by the phosphorylated PER-Short domain, we also performed kinase assays *in vitro* using the human CK1δ kinase domain and a PER2 PAS-Degron substrate in the presence of unphosphorylated and phosphorylated dPER-Short peptides (**Figure 7g**). CK1 phosphorylates only S589 in this peptide *in vitro* (Kivimae et al., 2008), although it is dependent upon NEMO kinase phosphorylation of S596 *in vivo* (Chiu et al., 2011). Here, we observed that peptides containing pS589, pS596, or both pS589/pS596 inhibited CK1 kinase activity similarly to the 3pFASP peptide (**Figure 7g**).

We then sought to characterize the structural basis for this inhibition with a crystal structure of CK1 in complex with the pS589 dPER-Short domain peptide (**Figure 7g-h**). This structure revealed a binding mode similar to the pFASP, where the substrate binding cleft was largely occluded by the peptide (**Figure 7h**) and anchored by the coordination of pS589 at anion binding Site 1 (**Figure 7i**). However, the activation loop of the kinase took on the rare ‘loop up’ conformation in this complex, which disrupts the second anion binding pocket (Philpott et al., 2020). This alternate conformation exposes a new channel that runs from the active site down towards the bottom of the kinase between the activation loop and helix-F that is bound by the dPER-Short peptide (**Figure 7h**), constricting at its narrowest point around dPER residue G593. We did not have density for residues after 595, including for pS596 in structures solved with the doubly phosphorylated peptide (data not shown). Therefore, although both mammalian and *Drosophila* PER peptides dock a critical CK1-dependent phosphoserine into anion binding Site 1, changes in peptide binding along the kinase active site suggest different mechanisms of recognition used to bind and inhibit the kinase.

## Discussion

The CK1 kinase family is defined by several highly conserved anion binding sites located around the substrate binding cleft that regulate kinase dynamics, substrate specificity, and temperature compensation of circadian rhythms (Lowrey et al., 2000; Philpott et al., 2020; Shinohara et al., 2017). It has long been proposed these anion binding sites mediate the recruitment of phosphorylated substrates to prime activity on the CK1 consensus motif pSxxS and/or bind the autophosphorylated tail of the kinase to inhibit its activity (Graves and Roach, 1995; Longenecker et al., 1996). Here, we show that CK1-dependent phosphorylation of key regulatory sites on its substrate PER near the kinase anchoring domain leads to feedback inhibition of the kinase through these conserved anion binding sites. Anchoring interactions can significantly enhance the kinetics of kinase activity on low to moderate affinity substrates nearby (Dyla et al., 2022), as was recently demonstrated for CK1 and its activity on low affinity, non-consensus phosphorylation sites on PER and FRQ (Marzoll et al., 2022). Inhibition of CK1 activity by phospho-PER is conserved in both mammals and *Drosophila* through anion binding Site 1, although other details of the PER-inhibited kinase complex differ by species. Notably, the loss of these CK1-dependent phosphosites in *Drosophila* (Konopka and Benzer, 1971), mice (Xu et al., 2007), and humans (Toh et al., 2001) shortens circadian period by several hours *in vivo*, demonstrating the functional importance of CK1 feedback inhibition on the molecular clock.

NMR spectroscopy allowed us to probe the kinetics and mechanism of the progressive, sequential phosphorylation of the FASP region in human PER proteins by CK1. The sequential nature of kinase activity here suggests that targeting the initial, rate-limiting priming step at the FASP region could be a powerful way to influence circadian rhythms. In fact, CK1 activity is influenced by other post-translational modifications on or around the FASP priming site, such as O-GlcNAcylation (Durgan et al., 2011; Kaasik et al., 2013) and acetylation (Levine et al., 2020), demonstrating how this region is poised to integrate different metabolic signaling inputs for control over the clock. Our data suggest that CK1 inhibition by the human PER2 pFASP region depends on similar spacing between successive phospho-serines on the FASP and the distance between anion coordination sites in the substrate binding region of CK1. Although the structures of inhibited CK1 illustrate a common binding mode for the first three phosphorylation sites of the FASP region, deletion of downstream phosphorylation sites in the U2OS *Per1* and *Per2* indel cell lines demonstrates that they also contribute to kinase inhibition and period control as suggested by our molecular dynamics simulations.

The CK1-dependent site in the *Drosophila* PER-Short domain (pS589) bound to the kinase at Site 1, similar to the priming serine of the human pFASP (pS662), although a change in the conformation of the kinase activation loop created a new peptide-binding channel that led the dPER peptide down toward the bottom of the kinase. We could not visualize density for the peptide after residue 595 in our structure, but substitution of dPER residues G593 or P597 also phenocopies the *per*^S^ mutation (S589N) (Baylies et al., 1992; Konopka and Benzer, 1971), suggesting that they also contribute to binding and feedback regulation of the kinase. Furthermore, we observed that phosphorylation of S596, either alone or in combination with S589, enhanced inhibition of CK1 by the PER-Short domain peptide *in vitro*, indicating that additional interactions between the C-terminal half of the PER-Short domain and CK1 may have been occluded by crystal packing.

Despite a high degree of similarity between the kinases, expressing mammalian CK1 variants in *Drosophila* does not always recapitulate their effects on circadian period (Fan et al., 2009; Sekine et al., 2008; Venkatesan et al., 2019; Xing et al., 2017; Xu et al., 2005). This could be due to differences in substrate identity or accessibility given changes in the molecular architecture of mammalian and *Drosophila* clocks (Patke et al., 2020). However, the differences we observed in the mechanism of CK1 feedback inhibition by human and *Drosophila* PER may contribute to these functional differences as well. In addition to this, allosteric links between anion binding Site 1 and Site 2 in human CK1 profoundly influence the structural dynamics of the activation loop (Philpott et al., 2020) and it is not yet known if the DBT kinase domain exhibits similar dynamics.

Molecular mechanisms that regulate the stable association between PER-CK1, PER stability, and repressive activity appear to be broadly conserved across eukaryotes. Notably, the CK1 binding motifs are conserved in PER homologs from mammals (Eide et al., 2005; Lee et al., 2004), *Drosophila* (Kim et al., 2007; Nawathean et al., 2007) and *C. elegans* (Jeon et al., 1999), and there is functional conservation with the two helical motifs (FCD1/2) that constitute the CK1 interaction domain in *Neurospora* FRQ (He et al., 2006; Liu et al., 2019). CK1 mediates the repressive activity of PER in the mammalian clock, where cryptochromes facilitate the recruitment of PER1/2-CK1 to the transcription factor CLOCK:BMAL1 in the nucleus, leading to its phosphorylation and displacement from DNA early in the repressive phase (Cao et al., 2021; Chiou et al., 2016). Similar mechanisms of displacement-type repression are also found in *Drosophila* (Patke et al., 2020) and *Neurospora* (He et al, 2006) circadian clocks. These mechanisms generally depend on CK1 activity and the strength of kinase anchoring (Cao et al., 2021; Liu et al., 2019), although DBT catalytic activity may not be required for displacement-type repression in *Drosophila* (Yu et al., 2009). Our data suggest a model whereby phosphorylation of key regulatory sites in PER close to the CK1BD results in feedback inhibition of CK1, thereby directly regulating the ability of CK1 to phosphorylate PER and possibly other substrates within the circadian clock.

It is not yet clear where in the cell and when in the ∼24-hr cycle of the molecular clock that regulation of CK1 activity by the PER pFASP/Per-Short domain occurs. In mammals, PER1/2 form stable complexes with CRY1/2 and CK1 in the cytoplasm (Aryal et al., 2017), although PER proteins remain largely hypophosphorylated until after nuclear entry (Beesley et al., 2020; Lee et al., 2001). PER phosphorylation is counterbalanced by phosphatases in mammals (Gallego et al., 2006b; Lee et al., 2011a; Schmutz et al., 2011) and *Drosophila* (Sathyanarayanan et al., 2004), so further work will be necessary to determine how kinase and phosphatase activity are integrated with other post-translational modifications to regulate FASP/PER-Short phosphorylation and influence feedback regulation of CK1 activity.

## Acknowledgements

We thank the staff at the 23-ID-D beamline at the Advanced Photon Source of the Argonne National Lab for their help with data collection. Funding for this work was provided by the US National Institutes of Health grants R01 GM107069 and R35 GM141849 (to C.L.P.), R01 GM131283 (to C.L.), R01 DK124068 (to J.C.C.), R01 GM31749 (to J.A.M.), and Singapore Ministry of Health grant MOH-000600 to D.M.V. S.R.H. was supported by NIH fellowship F32 GM133149.

## Author contributions

Conceptualization: J.M.P., S.R.H., C.L., and C.L.P.; Software: C.G.R. and J.A.M.; Investigation: J.M.P., A.F.M., J.P., K.L., S.R.H., R.N., D.H.S., R.R., and J.C.C.; Formal Analysis: J.M.P., A.M.F., S.R.H., R.N., Y.C., and S.T.; Writing – Original draft: J.M.P., C.G.R., C.L., and C.L.P.; Writing – Review & Editing: J.M.P., C.G.R., R.N., D.M.V., J.C.C., C.L., and C.L.P.; Visualization: J.M.P.; Supervision: J.A.M., D.M.V., J.C.C., C.L., and C.L.P.; Project Administration: C.L. and C.L.P.; Funding Acquisition: J.A.M., D.M.V., J.C.C., C.L., and C.L.P.

## Declaration of interests

The authors declare no competing interests.

## STAR Methods

### RESOURCE AVAILABILITY

#### Lead contact

*Further information and requests for resources and reagents should be directed to and will be fulfilled by the lead contact, Carrie L. Partch (*)

#### Materials availability

- Plasmids generated in this study are available upon request.
- U2OS reporter cell lines generated in this study are available upon request.

#### Data and code availability

- Data and code generated in this study are available upon request.
- NMR chemical shift assignments for the human FASP peptide have been deposited at BMRB and will be publicly available upon publication. The BMRB entry number is listed in the Key Resources Table.
- Crystal structures of human CK1δ in complex with phosphorylated PER peptides have been deposited to the PDB and will be publicly available upon publication. Accession codes are listed in the Key Resources Table.

### EXPERIMENTAL MODEL AND SUBJECT DETAILS

#### Cell lines

- The HEK293T cell line was purchased from ATCC (#CRL-3216).
- The *Drosophila* S2 cell line was purchased from Thermo Fisher Scientific (#R69007).
- The U2OS cell line was purchased from ATCC (#HTB-96).
  - Generation of *Per* KI cell lines is described in detail in Park et al, 2022.

#### *Per* mutant clones in *Per^Luc^* reporter cell lines

Heterozygous KI clones, H10 for *Per1^Luc^* and LH1 for *Per2^Luc^*, were used to generate mutations in *Per1* and *Per2*, respectively. These mRuby3-expressing reporter cells were infected with all-in-one CRISPR adenovirus (*Per2*-E17-S662) or transfected with all-in-one pAdTrack-Cas9-DEST plasmids expressing GFP as described previously (Jin et al., 2019). sgRNA sequence and selected clones are summarized in the table below. For clonal isolation of mutant cells, GFP-positive cells were sorted by FACS into 96-well plates, and these clones were further selected based on alterations in period and/or phase in bioluminescence rhythms. In each project, the majority of putative mutant clones showed a similar degree of period lengthening or shortening in bioluminescence screening. The clones used in this study were fully characterized by sequencing and immunoblotting (**Supplemental File 1a**.)

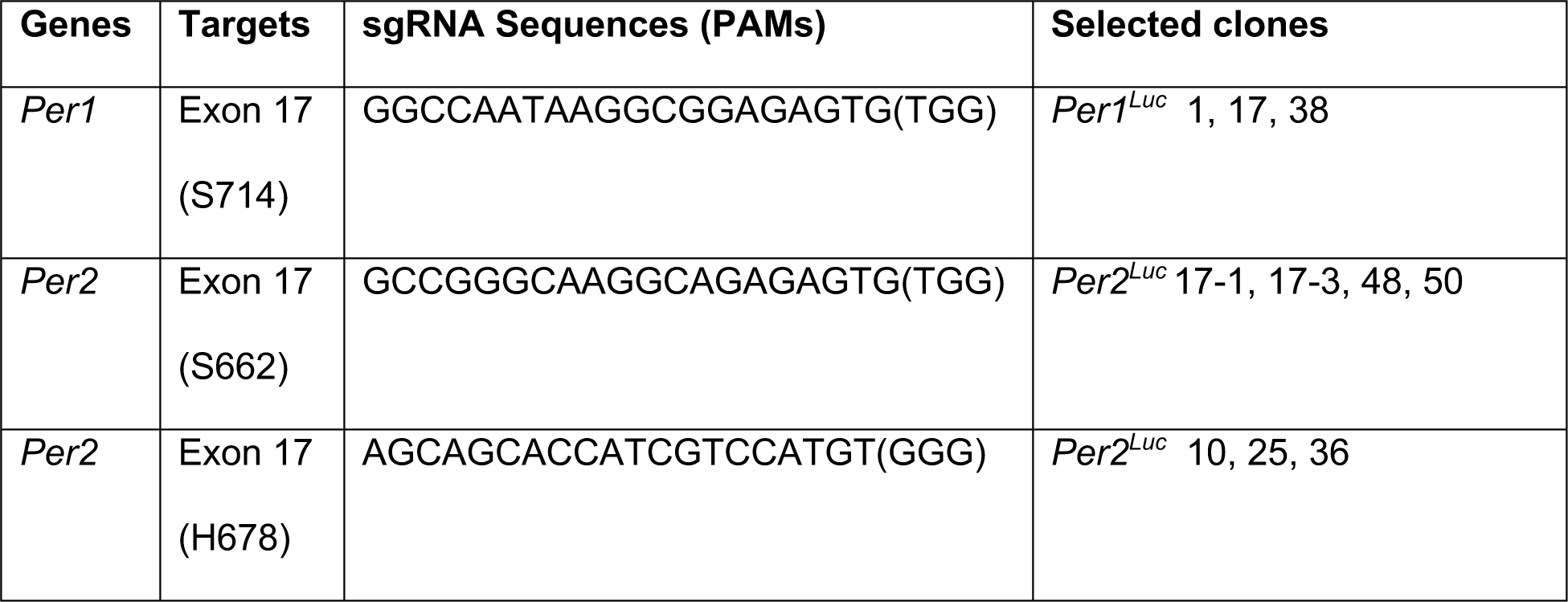

### METHOD DETAILS

#### Expression and purification of recombinant proteins

All proteins were expressed from a pET22-based vector in *Escherichia coli* BL21 (DE3) Rosetta2 cells (Sigma Aldrich) based on the Parallel vector series (Sheffield et al., 1999). The wild-type recombinant FASP peptide (residues 645–687) or PAS-Degron peptide (residues 475–505) were cloned from human PER2. All peptides were expressed downstream of an N-terminal TEV-cleavable His-NusA tag (HNXL). Human CK1δ catalytic domains (CK1δ ΔC, residues 1–317 for kinase assays and 1-294 for crystallography) were all expressed in BL21 (DE3) Rosetta2 cells (Sigma Aldrich) with a TEV-cleavable His-GST tag. Mutations were made using standard site-directed mutagenesis protocols and validated by sequencing. All proteins and peptides expressed from Parallel vectors have an N-terminal vector artifact (GAMDPEF) remaining after TEV cleavage and the peptides have a tryptophan and polybasic motif (WRKKK) following the vector artifact. Cells were grown in LB media (for natural abundance growths) or M9 minimal media with the appropriate stable isotopes, i.e., ^15^N/^13^C, for NMR as done before (Narasimamurthy et al., 2018) at 37°C until the O.D._600_ reached ∼0.8; expression was induced with 0.5 mM IPTG, and cultures were grown for approximately 16–20 hr more at 18°C.

For CK1δ kinase domain protein preps, cells were lysed in 50 mM Tris pH 7.5, 300 mM NaCl, 1 mM TCEP, and 5% glycerol using a high-pressure extruder (Avestin) or sonicator on ice (Fisher Scientific). HisGST-CK1δ ΔC fusion proteins were purified using Glutathione Sepharose 4B resin (GE Healthcare) using standard approaches and eluted from the resin using Phosphate Buffered Saline with 25 mM reduced glutathione. His-TEV protease was added to cleave the His-GST tag from CK1δ ΔC at 4°C overnight. Cleaved CK1δ ΔC was further purified away from His-GST and His-TEV using Ni-NTA resin (Qiagen) and subsequent size exclusion chromatography on a HiLoad 16/600 Superdex 75 prep grade column (GE Healthcare) in 50 mM Tris pH 7.5, 200 mM NaCl, 5 mM BME, 1 mM EDTA, and 0.05% Tween 20. Purified CK1δ ΔC proteins used for *in vitro* kinase assays were buffer exchanged into storage buffer (50 mM Tris pH 7.5, 100 mM NaCl, 1 mM TCEP, 1 mM EDTA, and 10% glycerol) using an Amicron Ultra centrifugal filter (Millipore) and frozen as small aliquots in liquid nitrogen for storage at -80°C.

For PER2 peptide preps, cells were lysed in a buffer containing 50 mM Tris pH 7.5, 500 mM NaCl, 2 mM TCEP, 5% glycerol and 25 mM imidazole using a high-pressure extruder (Avestin) or sonicator on ice (Fisher Scientific). His-NusA-FASP or His-NusA-PAS-Degron fusion proteins were purified using Ni-NTA resin using standard approaches and eluted from the resin using 50 mM Tris pH 7.5, 500 mM NaCl, 2 mM TCEP, 5% glycerol and 250 mM imidazole. His-TEV protease was added to cleave the His-NusA tag from the PER2 peptides at 4°C overnight. The cleavage reaction was subsequently concentrated and desalted into low imidazole lysis buffer using a HiPrep 26/10 Desalting column. Peptides were purified away from His-NusA and His-TEV using Ni-NTA resin with 50 mM Tris pH 7.5, 500 mM NaCl, 2 mM TCEP, 5% glycerol and 25 mM imidazole. Peptides were purified by size exclusion chromatography on a HiLoad 16/600 Superdex 75 prep grade column, using NMR buffer (25 mM MES pH 6.0, 50 mM NaCl, 2 mM TCEP, 10 mM MgCl2) or 1x kinase buffer (25 mM Tris pH 7.5, 100 mM NaCl, 10 mM MgCl2, and 2 mM TCEP) for NMR or ADP-Glo kinase assays, respectively.

#### NMR kinase assays

NMR spectra were collected on a Varian INOVA 600 MHz or a Bruker 800 MHz spectrometer equipped with a ^1^H, ^13^C, ^15^N triple resonance z-axis pulsed-field-gradient cryoprobe. Spectra were processed using NMRPipe (Delaglio et al., 1995) and analyzed using CCPNmr Analysis (Vranken et al., 2005). Backbone resonance assignments were made using standard BioPack triple resonance experiments (HNCACB, CBCA (CO)NH, HNCO, HN (CA)CO and HSQC) collected using non-uniform sampling on a sample of 0.5 mM ^13^C, ^15^N-labeled FASP in NMR buffer with 10% D2O. Non-uniform sampling reconstructions were performed using software developed and provided by the Wagner lab (Delaglio et al., 1995). NMR kinase reactions were performed at 25°C with 0.2 mM ^15^N-FASP, 2.5 mM ATP and 1 µM CK1δ ΔC (WT or K224D). SOFAST HMQC spectra (total data acquisition = 6 min) were collected at the indicated intervals for 3 hr and relative peak volumes were calculated and normalized as described previously (Narasimamurthy et al., 2018). For PER1 FASP peptide kinase assays, samples were prepared as above and incubated for 2 hrs and then quenched with 20 mM EDTA and then HSQC spectra were collected. Data analysis was performed using Prism (GraphPad), with data fit to either a one-phase exponential or linear regression.

#### Kinetic modeling

Mathematica 11.0 (Wolfram Research) was used to model the 5-step ordered distributive kinetic model for sequential FASP phosphorylation (**Figure 2a-c**). Mathematica code is provided in the supplementary information (**Supplementary File 1c)**.

#### ADP-Glo kinase assays (substrate titrations)

Kinase reactions were performed on the indicated recombinant peptides (FASP WT or Alanine mutants) using the ADP-Glo kinase assay kit (Promega) according to manufacturer’s instructions. All reactions were performed in 30 μL volumes using 1x kinase buffer (25 mM Tris pH 7.5, 100 mM NaCl, 10 mM MgCl2, and 2 mM TCEP) supplemented with ATP and substrate peptides. To determine apparent kinetic parameters K_M_ and V_max_, duplicate reactions with 100 µM ATP and 0.2 µM CK1δ ΔC kinase were incubated in 1x kinase buffer at room temperature for 1 hr with the indicated amount of substrate peptide (and repeated for n = 3 independent assays). 5 µL aliquots were taken and quenched with ADP-Glo reagent after the 1 hr incubation, and Luminescence measurements were taken at room temperature with a SYNERGY2 microplate reader (BioTek) in 384-well microplates. Linearity of the 1 hr reaction rate was determined by performing larger reactions (50 µL) with CK1δ ΔC and quenching at discrete time points (data not shown). Data analysis was performed using Excel (Microsoft) or Prism (GraphPad).

#### ADP-Glo kinase assays (peptide inhibition)

Kinase reactions were performed on the recombinant PAS-Degron peptide (**Supplementary Figure 2b**) using the ADP-Glo kinase assay kit (Promega) according to manufacturer’s instructions. Synthetic peptide inhibitors were solubilized in 1x kinase buffer (25mM Tris pH 7.5, 100 mM NaCl, 10 mM MgCl_2_, and 2 mM TCEP) at a concentration of 10 mM. All kinase reactions were performed in 30 μL reactions supplemented with ATP, PAS-Degron substrate, and synthetic peptide inhibitors. Duplicate reactions (repeated for n = 3 independent assays) with 100 µM ATP, 0.2 µM CK1δ ΔC kinase, and 100 µM PAS-Degron substrate were incubated in 1x kinase buffer at room temperature for 1 hr in the presence of increasing amounts of synthetic peptide inhibitors (phosphorylated FASP or dPER peptides) as indicated. 5 µL aliquots were quenched with ADP-Glo reagent after the 1 hr incubation, and luminescence measurements were taken at room temperature with a SYNERGY2 microplate reader (BioTek) in 384-well microplates. Data analysis was performed using Excel (Microsoft) or Prism (GraphPad).

#### Full-length PER2 kinase assay (peptide inhibition)

HEK293 cells transfected with Myc-PER2 plasmid were lysed in cell lysis buffer (50 mM Tris-HCl pH 8.0, 150 mM NaCl, 1% Nonidet P-40, and 0.5% deoxycholic acid containing Complete protease inhibitors (Roche) and PhosStop phosphatase inhibitors (Roche)). 300 μg of the protein lysate was added with 3 μg of anti-Myc antibody (9E10) and allowed to rotate at 4°C for 1 hr, which was followed by addition of Protein A/G magnetic beads (Thermo Scientific) and rotation at 4°C for 1hr. Then the beads were collected and washed 3x with lysis buffer. Beads were then collected and treated with FastAP alkaline phosphatase (Thermo Scientific) for 30 min at 37°C and washed 3x with lysis buffer and 2x with kinase assay buffer (25 mM Tris pH 7.5, 5 mM beta glycerophosphate, 2 mM DTT and 0.1 mM sodium orthovanadate). CK1δ ΔC (200 ng) was incubated with either 1 mM of non-phospho (NP) FASP peptide RKKK(mouse PER2 residue 642)TEVSAHLSSLTLPGKAESVVSLTSQ (Narasimamurthy et al., 2018) or the human PER2 4pFASP peptide GKAEpSVApSLTpSQCpSYA for 15 min at 25°C. The beads were split into three samples with kinase assay buffer containing 10 mM of magnesium chloride and 200 μM of ATP. The first portion was left untreated while the second and third were incubated with CK1δ-ΔC (200 ng) that had been pre-incubated with NP FASP or 4pFASP peptide, respectively, and incubated for 60 min at 25°C. The beads were collected by centrifugation and protein was eluted by adding protein loading dye and analyzed by SDS-PAGE gel for Western blotting with anti-mouse PER2 pSer 659 antibody as previously described (Narasimamurthy et al., 2018).

#### Real-time PER2::LUC half-life measurement

1 μg of the indicated human PER2::LUC expression plasmids (under a PGK promoter) were transiently transfected alone or with 100 ng myc-CK1ε (under a CMV promoter) in 35 mm dishes of HEK293T cells in MOPS-buffered high glucose media supplemented with D-luciferin (8.3g/L DMEM, 0.35 mg/mL sodium bicarbonate, 5 mg/mL glucose, 0.02 M MOPS, 100 U/mL penicillin, 100 µg/mL streptomycin, and 100 mM D-luciferin). Dishes were sealed with 40 mm cover glasses and vacuum grease (Sigma Aldrich), and then placed in a LumiCycle 32 (Actimetrics) at 37°C. 24 hrs post transfection, 40 μg/mL cycloheximide (Sigma) was added per 35 mm dish and luminescence recording was initiated. Luminescence data were used to calculate PER2::LUC half-life in Prism (GraphPad) using one-phase decay algorithm as described previously (Zhou et al., 2015), beginning from the point of cycloheximide addition to the plateau at minimum luciferase activity (n ≥ 4).

#### Co-immunoprecipitation assays

Human myc-PER2, FLAG-CK1δ and FLAG-β-TrCP expression plasmids (1.5 μg, 1.5 μg WT or 0.75 μg K38A, and 3 μg of plasmid DNA, respectively) were co-transfected in 60 mm dishes of HEK293T cells in DMEM supplemented with 1% penicllin/streptomycin and 10% HyClone FetalClone II FBS (Fisher Science). The proteasome was inhibited 16 hrs prior to harvest with 10 μM MG132 (Sigma Aldrich) and cells were harvested 72 hrs post transfection. Cells were lysed in 350 μL mammalian cell lysis buffer (20 mM Tris pH 7.5, 150 mM NaCl, 1mM TCEP, 1% NP-40) supplemented with EDTA-free protease inhibitors (Pierce) and phosphatase inhibitors (1 mM NaF, 1 mM β-glycerophosphate and 1 mM Na_3_VO_4_). 30 μL input samples were prepared with 2x SDS sample buffer, and the remaining lysate was incubated with 30 μL of anti-Myc agarose slurry (cat. #sc-40 AC, Santa Cruz Biotechnology) and tumbled overnight at 4°C. Bound samples were washed 3x with 400 μL of IP wash buffer (20 mM Tris pH 7.5, 150 mM NaCl) and 6x SDS sample buffer. All samples were briefly boiled at 95°C and resolved by 7.5% polyacrylamide-SDS gel electrophoresis (PAGE) and transferred to nitrocellulose membrane via Trans-Blot Turbo transfer system (Bio-Rad). Membranes were incubated in TBST blocking buffer (20 mM Tris pH 7.5, 150 mM NaCl, 0.1% TWEEN 20 and 5% (weight/vol, w/v) Marvel dried skimmed milk) for 1 hr and then incubated with antibody (1:2000 α-Myc HRP (cat. #sc-40 HRP), Santa Cruz Biotechnology) or 1/1000 α-OctA HRP (cat. #sc-166355 HRP, Santa Cruz Biotechnology)) in blocking buffer at 4°C overnight. Membranes were washed the next day 3x with TBST and chemiluminescence detection was performed using Immobilon reagent (Millipore) and imaged with ChemiDoc (Bio-Rad). Representative blot shown (n = 3).

#### Crystallization and structure determination

All peptides were solubilized in a solution of 0.15 M Succinic Acid pH 5.5 and 20% (w/v) PEG 3350 and soaked overnight into crystals of human CK1δ (1-294) that were crystallized using the hanging drop vapor diffusion method as follows: the 4pFASP peptide was added to CK1δ crystals that were crystallized in 0.13 M Succinic Acid pH 5.5 and 27% (w/v) PEG 3350; the 3pFASP peptide was added to CK1δ crystals that were crystallized in 0.1 M Succinic Acid pH 5.5 and 15% (w/v) PEG 3350; the 2pFASP peptide was added to CK1δ crystals that were crystallized in 0.15 M Succinic Acid pH 5.5 and 22% (w/v) PEG 3350; and the pS589 peptide was added to CK1δ crystals that were crystallized in 0.16 M Succinic Acid pH 5.5 and 23% PEG (w/v) 3350. The crystals were looped and briefly soaked in a drop of cryo-preservation solution (80% peptide solution, 20% glycerol) and then flash-cooled in liquid nitrogen for X-ray diffraction data collection. Data sets were collected at the 23-ID-D beamline at the Advanced Photon Source (APS) at the Argonne National Laboratory. Data were indexed, integrated and merged using the CCP4 software suite (Winn et al., 2011). Structures were determined by molecular replacement with Phaser MR (McCoy et al., 2007) using the apo structure of wild-type CK1δ ΔC (PDB: 6PXO). Model building was performed with Coot (Emsley et al., 2010) and structure refinement was performed with PHENIX (Adams et al., 2011). All structural models and alignments were generated using PyMOL Molecular Graphics System 2.0 (Schrödinger). X-ray crystallography data collection and refinement statistics are provided in supplementary information (**Table 1**).

#### pLogogram analysis

The Comparative Site Search function of PhosphoSitePlus (v6.6.0.4) (Hornbeck, 2015) was used to obtain a list of known CK1δ substrates (**Supplementary File 1b**). The substrates were all limited to 15 residues in length with the center residue being the site of phosphorylation. The Sequence Logo Analysis tool from PhosphoSitePlus was used on the resulting dataset to generate a pLogogram indicating the relative frequency of amino acid types in positions flanking the site of phosphorylation.

#### Molecular dynamics

##### Molecular modeling

To build the molecular model of 5pFASP-CK1, we started from the 3pFASP x-ray structure and then added E661 and residues 669 to 675 with Maestro (Schrodinger Release 2020-3) in a linear conformation. To avoid artificial interactions arising from the short size of the peptide, we capped the N- and C-terminal residues with acetyl (ACE) and N-methyl amine (NME) groups, respectively. Hydrogens were added with PrepWizard module, and a restrained minimization was to remove potential clashes between the modeled FASP peptide and the enzyme.

##### *System* preparation

CK1-5pFASP complex was solvated in a pre-equilibrated TIP3P (Jorgensen et al., 1983) water box with at least 15 Å between the protein and the box boundaries. The net charge of the system was neutralized with a Na^+^ ion. Parameters for protein residues, capping groups, and Na+ were obtained from ff14SB forcefield (Maier et al., 2015).

##### Equilibration

Minimization and equilibration were performed with AMBER16 (Case, 2016) using the following protocol: (i) 2000 steps of energy minimization with 500 kcal mol^-1^ Å^-1^ position restraints on all protein atoms; (ii) 5000 steps of energy minimization with 500 kcal mol^-1^ Å^-1^ position restraints on all CK1 atoms and on the backbone atoms of FASP; (iii) 5000 steps of energy minimization with 500 kcal mol^-1^ Å^-1^ position restraints on all backbone atoms; (iv) 5000 steps of energy minimization with 500 kcal mol^-1^ Å^-1^ position restraints on CK1 backbone atoms; (v) 5000 steps of energy minimization without any position restraints; (vi) 50 ps of NVT simulations, with gradual heating of the system to a final temperature of 300 K and 10 kcal mol^-1^ Å^-1^ position restraints on protein atoms; (vii) 1 ns of NPT simulation with 100 kcal mol^-1^ Å^-1^ position restraints on protein atoms; and (viii) 1 ns of NPT simulation to equilibrate the density (or final volume of the simulation box).

##### Simulations

Gaussian accelerated MD simulations (GaMD) were performed in the NVT regime, with a time step of 2 fs. The PME method was used to calculate electrostatic interactions with periodic boundary conditions (Darden et al., 1993). To accelerate sampling of the conformational space, we used boost parameters as described in (Miao et al., 2015). All systems had a threshold energy V = V_max_ and were subjected to a dual boost acceleration of both dihedral and total potential energies. To optimize the acceleration parameters, we first ran 2 ns of conventional MD simulation (without boost potentials) during which V_min_, V_max_, V_avg_, and σ_avg_ were recorded and used to calculate boost potentials as previously detailed (Miao et al., 2015). These potentials were employed to start 50 ns of preparatory GaMD simulations, during which the boost statistics and boost potentials were updated until the maximum acceleration was achieved. The maximum acceleration was constrained by setting the upper limit of the standard deviation of the total boost potential to be ≤ 6 kcal/mol. Starting from the same equilibrated structure, we launched 10 independent GaMD simulations with different initial velocities. Each simulation ran for 100 ns, totalizing 1 μs of sampling time.

#### Bioluminescence recording of *Per^Luc^* reporter cell lines

Cells were plated into 24-well plates or 35 mm dishes to be approximately 90% confluent 24 hours prior to the start of the experiment. Immediately before the start of the experiment, cells were given a two-hour serum shock with 50% horse serum in DMEM or 10 μM forskolin (Sigma Aldrich) in DMEM (Fig S5), washed with phosphate-buffered saline (PBS) and fresh DMEM supplemented with 1% FBS, 7.5 mM sodium bicarbonate, 10 mM HEPES, 25 U/mL penicillin, 25 μg/mL streptomycin, and 0.1 mM luciferin. The plates were sealed with cellophane tape and the dishes with a 40 mm cover glass and vacuum grease before placing them into a Lumicycle 32 or 96 (Actimetrics). For all bioluminescence experiments, the results were reproduced in at least two independent experiments. Real-time levels, period, and phase of the bioluminescence rhythms were evaluated using the Lumicycle software (Actimetrics). Student’s t-test was used to compare data from WT and mutant cells.

##### Circadian sampling and drug treatments in *Per^Luc^* reporter cell lines

To measure PER rhythms in WT and mutant cells, cells were seeded in 60 mm dishes to be 90% confluent 24 hours prior to the experiment. These cells were treated with 50% horse serum in DMEM for 2 hrs and harvested at the indicated times. For cycloheximide (CHX) treatment, 8 μg/mL CHX (Sigma Aldrich) was added to cells and cells were collected at specified times after the treatment. For CHX washout experiments, cells were treated with CHX for 8 hrs followed by the normal DMEM, and cells were harvested at the indicated times after the CHX treatment.

#### Immunoblotting of *Per^Luc^* reporter cell lines

Cells were harvested from 60 mm dishes and flash-frozen on dry ice. Protein extraction and immunoblotting were performed as previously described (D’Alessandro et al., 2015). Briefly, cells were homogenized at 4°C in 70 μL extraction buffer (EB) (0.4 M NaCl, 20 mM HEPES pH 7.5, 1 mM EDTA, 5 mM NaF, 1 mM dithiothreitol, 0.3% Triton X-100, 5% glycerol, 0.25 mM phenylmethylsulfonyl fluoride (PMSF), 10 mg/mL aprotinin, 5 mg/mL leupeptin, 1 mg/mL pepstatin A). Homogenates were cleared by centrifugation for 12 min at 12,000 g at 4°C. Supernatants were mixed with 2x SDS sample buffer and boiled. Proteins were separated by electrophoresis through SDS polyacrylamide gels and then transferred to nitrocellulose membranes. Membranes were blocked with 5% (w/v) non-fat dry milk in TBS-0.05% Tween-20 (TBST), incubated with primary antibodies (GP62 for PER1 and GP49 for PER2 (Jin et al., 2019)) overnight followed by incubation with secondary antibodies for 1 h. The blots were developed using the WestFemto enhanced chemiluminescence substrate (Thermo Fisher Scientific).

#### *Drosophila* S2 cell experiments

The plasmids pMT-*dbt*-V5, pAc-3XFLAG-His-d*per/*Tev100-6Xc-myc (Chiu et al., 2008), pAc-d*per* WT (aa1-1224)-V5, pAc-d*per* S596A (aa1-1224)-V5 (Chiu et al., 2011), pAc-d*per* WT (aa560-1034)-V5 (Ko et al., 2010) were previously described. pAc-d*per* S596A (aa560-1034)-V5 was generated via PCR mutagenesis using pAc-d*per* WT (aa560-1034)-V5 as a template as described in (Chiu et al., 2011) and the QuikChange II site-directed mutagenesis kit (Agilent). *Drosophila* S2 cells and *Schneider’s Drosophila* medium were obtained from Life Technologies (Thermo Fisher Scientific). S2 cells were seeded at 1 x 10^6^ cells/mL in a 6-well plate and transfected using Effectene (Qiagen). S2 cells were co-transfected with 0.2 μg of pMT-*dbt*-V5, 0.8 μg of pAc-3X-FLAG-His-d*per/*Tev100-6Xc-myc, and varying amounts of d*per*-V5 plasmids as indicated. Expression of *dbt* was induced with 500 μM CuSO_4_ 36 hours after transfection. Cells were harvested at the indicated time points after kinase induction and extracted with 100-150 µL EB2 (20 mM Hepes pH 7.5, 100 mM KCl, 5% glycerol, 5 mM EDTA, 1 mM DTT, 0.1% Triton X-100, 25 mM NaF, 0.5 mM PMSF). 1 µL of AcTEV protease (Thermo Fisher Scientific) was added to protein extracts and incubated 16 hours at 4°C. Following TEV cleavage, protein concentration was measured using Pierce Coomassie Plus Assay Reagents (Thermo Fisher Scientific). 2X SDS sample buffer was added and the mixture boiled at 95°C for 5 minutes. Equal amounts of proteins were resolved by 16% polyacrylamide-SDS gel electrophoresis (PAGE) and transferred to nitrocellulose membrane (Bio-Rad) using a Semi-Dry Transfer Cell (Bio-Rad). Membranes were incubated in 5% Blocking Buffer (Bio-Rad) for 40 minutes at room temperature, and then incubated with anti-FLAG (cat. # F1804, Millipore Sigma) at 1:7000 for 16-20 hours at room temperature. Blots were washed with 1X TBST for 1 hour, incubated with anti-mouse IgG HRP (cat. #12-349, Millipore Sigma) at 1:2000 for 1 hour. After washing, chemiluminescence detection was performed using Clarity ECL reagent in combination with the ChemiDoc imager (Bio-Rad). ImageJ Version 2.0.0-rc-67/1.52d (NIH) was used to quantify hypo-phosphorylated PER and nonspecific signals (indicated by asterisks). PER signal was normalized to nonspecific signal and the data were scaled with normalized PER signal at 0 hr to a value of 1. Two-way ANOVA with Sidak multiple comparisons test was performed using Prism (GraphPad).

### QUANTIFICATION AND STATISTICAL ANALYSIS

All statistical analyses were done using Prism (Graphpad). P-values were calculated using Students t-test, one-way ANOVA with Tukey’s multiple comparisons test, or two-way ANOVA with Sidak multiple comparisons test as indicated in different figures. In all figures, * indicates p<0.05, **p<0.01, ***p<0.001, ****p<0.0001; ns, not significant.

#### Supplemental File Titles and Legends

Supplementary Table 1. Sequencing details for mutant *PER^Luc^* U2OS cell lines.

Supplementary Table 2. PhosphositePlus kinase substrate data set used for pLogogram analysis.

Supplementary Table 3. Kinetic modeling of ordered distributive mechanism using Mathematica.

## Notes

### Competing Interest Statement

The authors have declared no competing interest.

