## Supplemental Figures for "PERIOD phosphorylation leads to feedback inhibition of CK1 activity to control circadian period"

Supplemental Figure 1

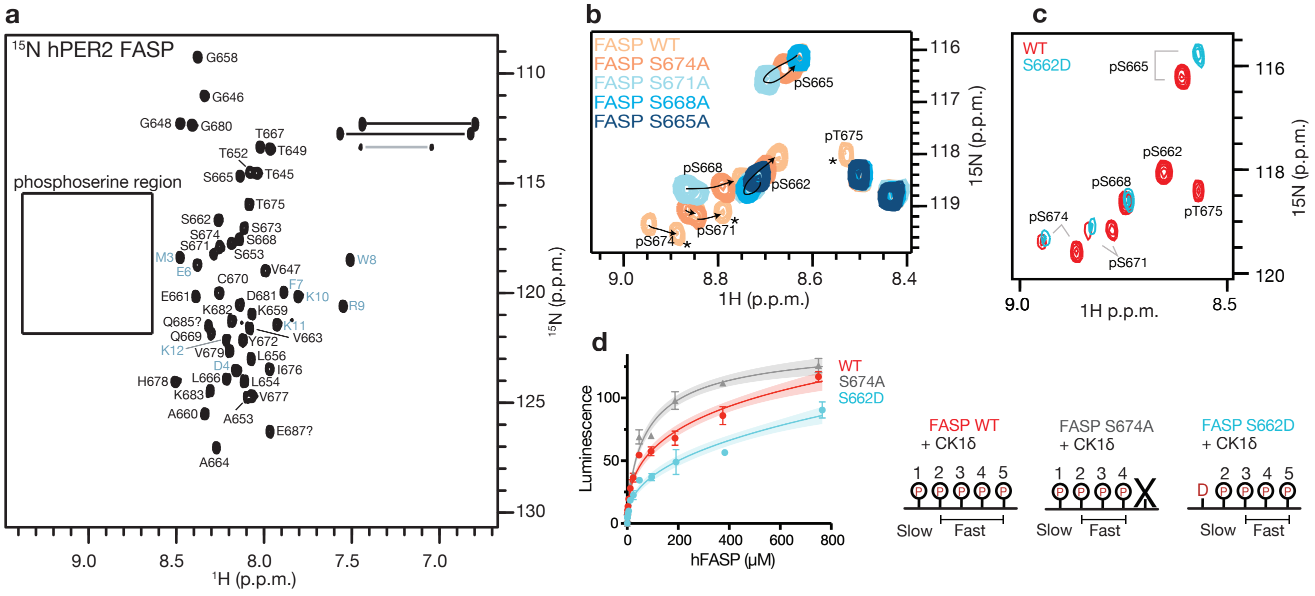

### Supplemental Figure 2

**a**

| FASP Substrate | Michaelis $K_m$ (uM) | Michaelis $V_{max}$ (pmol ATP/sec) | Substrate Inhibition $K_m$ (uM) | Substrate Inhibition $V_{max}$ (pmol ATP/sec) | Michaelis Fit $R^2$ | Substrate Inhibition Fit $R^2$ | Null Hypothesis (Michaelis Fit) P value |
|---|---|---|---|---|---|---|---|
| WT | 17.74 ± 0.45 | 0.266 ± 0.001 | 32.29 ± 2.05 | 0.355 ± 0.012 | 0.957 | 0.994 | <0.0001 |
| S674A | 26.61 ± 2.34 | 0.426 ± 0.011 | 50.58 ± 6.96 | 0.601 ± 0.006 | 0.965 | 0.992 | <0.0001 |
| S671A | 75.31 ± 0.61 | 0.610 ± 0.007 | 93.68 ± 0.84 | 0.692 ± 0.001 | 0.997 | 0.998 | 0.0027 |
| S668A | 102.90 ± 15.02 | 0.712 ± 0.026 | 118.20 ± 6.58 | 0.779 ± 0.070 | 0.997 | 0.997 | 0.7210 |

**b**

β-TrCP motif

PAS-Degron: GAMDPEFWRKKKPHSGSSGYGSLGSNGSHEHLMSQTSSSDSNG

S480 S484

**c**

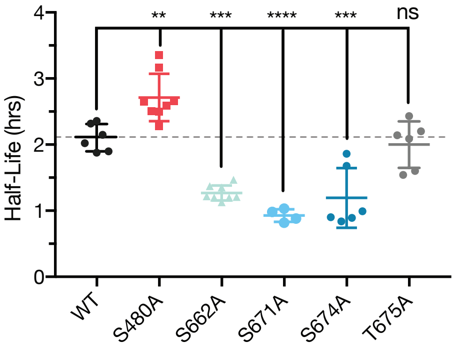


**d**

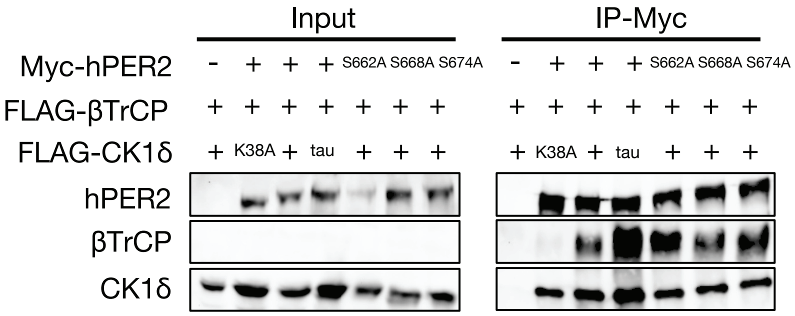

Supplemental Figure 3

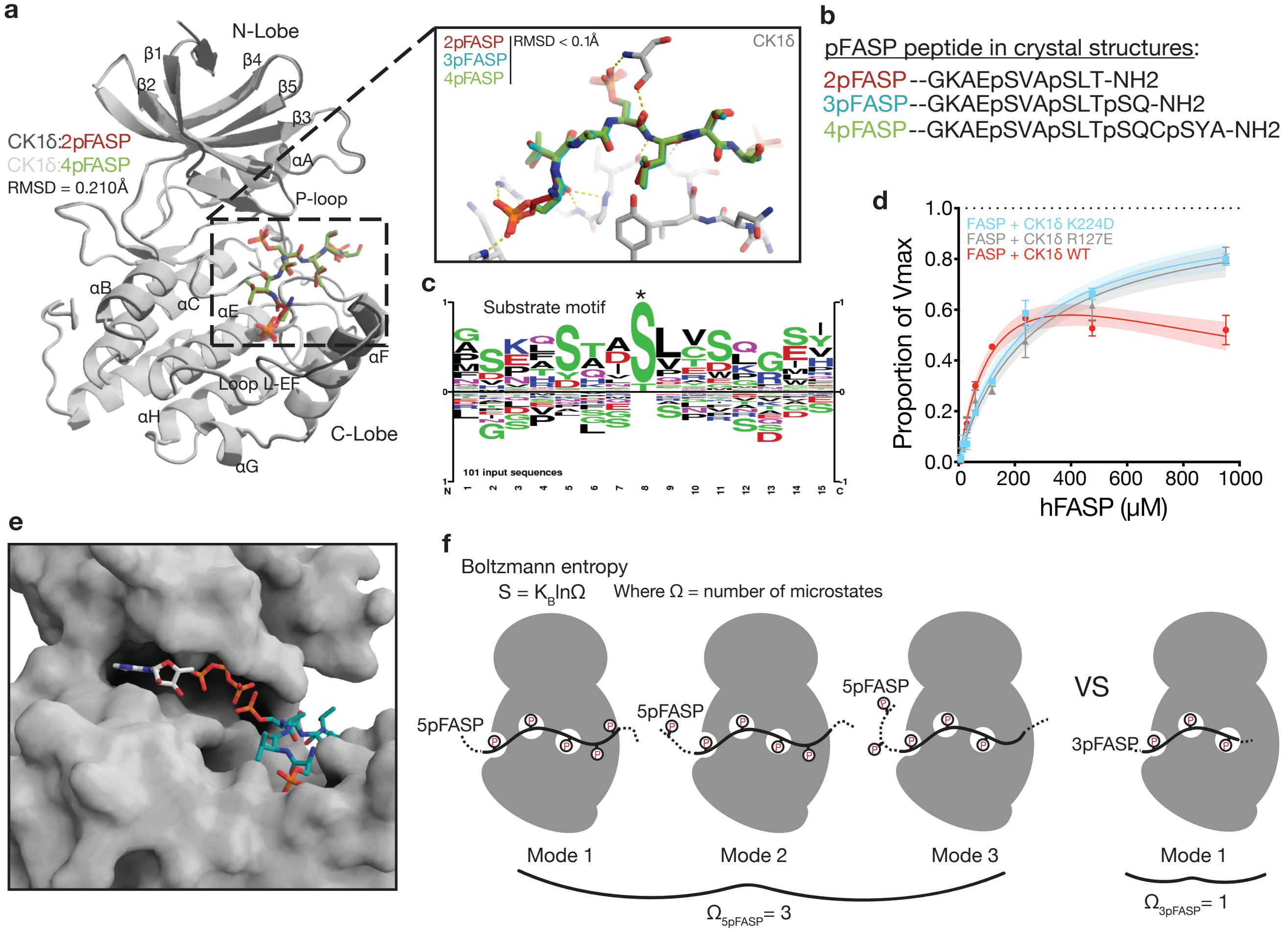

### Supplemental Figure 4

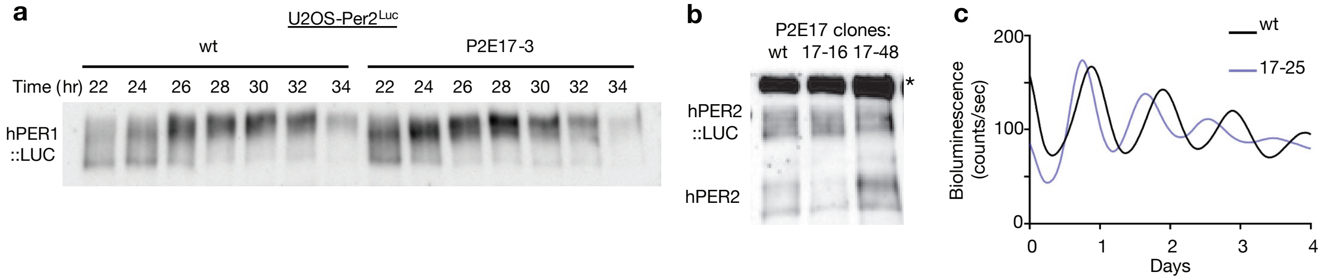

### Supplemental Figure 5

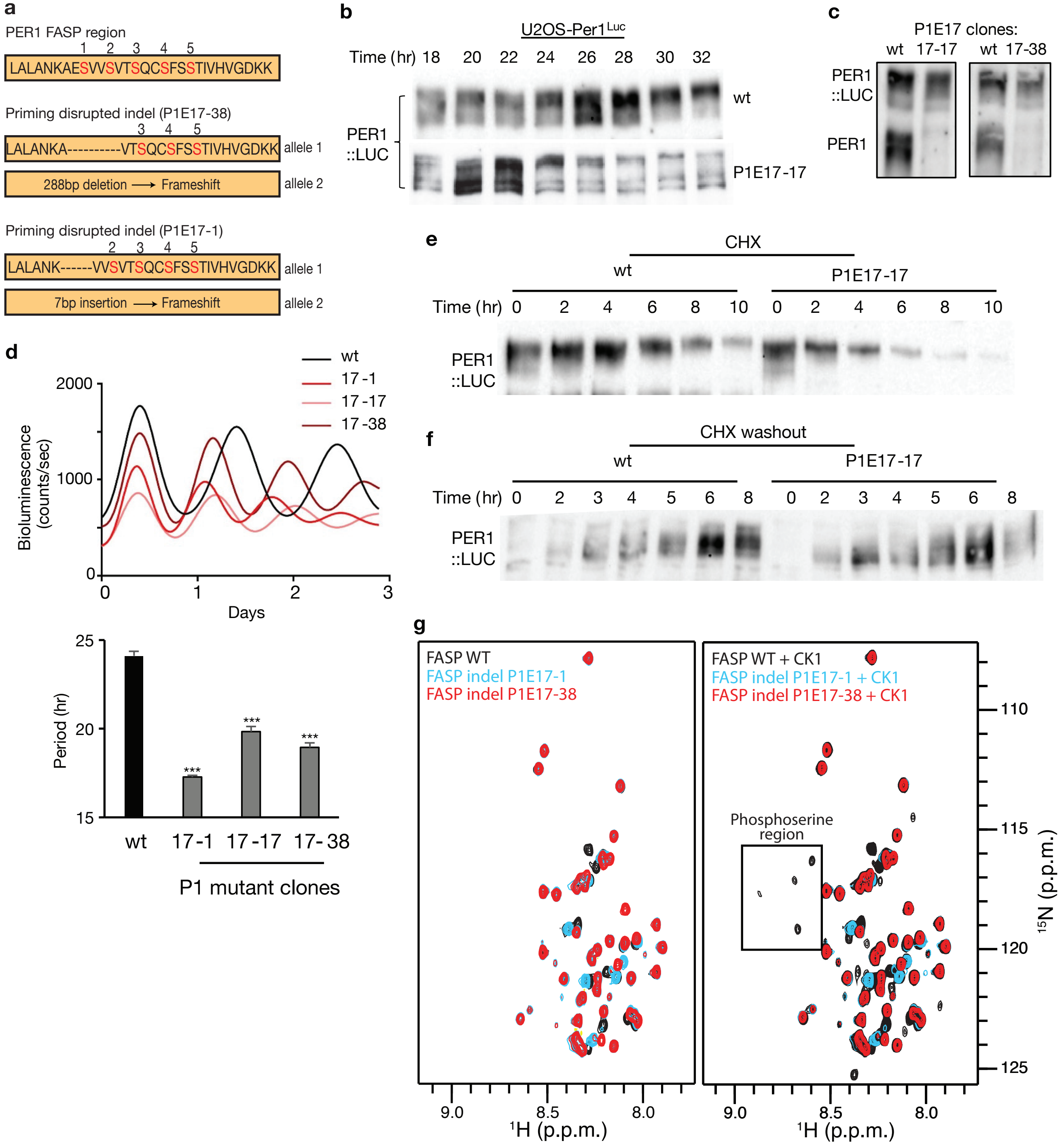

**Supplementary Figure 1 (relates to Figure 1 and Figure 2).**

**a**, $^{15}N/^{1}H$ HSQC of hPER2 FASP peptide with assignments. Boxed region indicates area within the spectra where phosphoserines appear. **b**, Overlaid $^{15}N/^{1}H$ HSQC spectra of FASP Ser/Ala mutant peptides. Arrows indicate movement of unique chemical shifts corresponding to extent of FASP phosphorylation. Asterisk indicates peaks with chemical shift due to phosphorylation of T675. **c**, Zoom into phospho-serine region of $^{15}N/^{1}H$ HSQC of hPER2 FASP peptide (WT) overlaid with the S662D phosphomimetic mutant after a 3 hr incubation with CK1δ and ATP at 25 ºC. **d**, ADP-Glo kinase assay with titration of FASP mutant peptides with mean and SD from 2 replicates, representative of n = 3 independent assays. Shaded area indicates 95% C.I. of the fit. Schematic represents the different combination of non-consensus and consensus-based phosphorylation events for each of the mutants. Note how the S674A and S662D mutants have an equivalent number of consensus-based phosphorylation sites.

**Supplementary Figure 2 (relates to Figure 3).**

**a**, Table of kinetic analysis for ADP-Glo assay in Figure 3, panel **f**. **b**, The sequence of the PAS-Degron peptide used for kinase substrate in pFASP inhibition assays (Figure 3, panel **h**) in black (blue, non-native amino acids). **c**, Quantification of real-time luciferase traces treated with cycloheximide 48 hrs after co-transfection of HEK293T cells with hPER2-LUC and CK1δ expression plasmids ($n \geq 4$ with SD). Significance assessed relative to WT with ordinary one-way ANOVA: **, $p < 0.01$; ***, $p < 0.001$; ****; p <

0.0001; ns, not significant. **d**, Co-immunoprecipitation assay from transiently transfected HEK293T cells assessing CK1δ-mediated β-TrCP recruitment to PER2 (n = 3).

**Supplementary Figure 3 (relates to Figure 4).**

**a**, Cartoon representation of CK1δ:2pFASP and CK1δ:4pFASP aligned structures to show overall similarity (RMSD indicated). Boxed region represents activation loop zoom. 2pFASP, 3pFASP, and 4pFASP aligned based on overall structural alignment of CK1δ:2pFASP and CK1δ:4pFASP complexes with CK1δ:3pFASP, with RMSD of peptides indicated. **b**, Synthetic peptides used for soaking into CK1δ crystals and ADP-Glo inhibition assays. **c**, Substrate motif analysis (pLogogram) generated with dataset of 101 known CK1δ substrates from PhosphositePlus. Relative size of residue-letter indicates relative frequency of that residue at that position. Asterisk indicates site of phosphorylation. **d**, ADP-Glo kinase assay with titration of hFASP peptide showing that anion binding site 1 (K224D) and anion binding site 2 (R127E) contribute to pFASP inhibition. Data (mean and SD of 2 replicates) are normalized to Vmax calculated from the preferred model fit (Michaelis or Substrate Inhibition). Shaded area indicates 95% C.I. of the fit. **e**, Zoom in of CK1δ:3pFASP active site. ATP is shown in active site based on alignment of the CK1δ:3pFASP catalytic domain with an ATP bound structure of CK1δ, 1CSN. **f**, Schematic representation of entropic differences between 3pFASP and 5pFASP binding modes. The higher number of accessible microstates in 5pFASP could contribute to the observed increase in inhibition of kinase activity.

**Supplementary Figure 4 (relates to Figure 6).**

**a**, PER1 in the PER2-LUC mutant clone 17-3 shows phase-advanced rhythms in abundance and phosphorylation (n = 2).

**b**. Two mutant clones 17-16 and 17-48 are shown. * non-specific band. **c**, Clone P2E17-25 showed shorter period bioluminescence rhythms.

**Supplementary Figure 5 (relates to Figure 6).**

**a**, Schematic representation of select in-frame deletions within the PER1 FASP region. **b**, Targeting S714 with CRISPR produced PER1 mutants with rhythms in protein abundance that peak earlier relative to wt. The PER1 blot for 17-17 is shown. **c**, Representative immunoblots for mutants and wt PER1. **d**, Real-time bioluminescence traces of *Per1* mutants and quantification of periods. Each period represents the mean and SD calculated from n = 3 cultures, compared to wt cells with an unpaired t-test. **e**, Degradation of PER1 wt and clone 17-17 after cycloheximide treatment (n = 2). **f**, Phosphorylation of *de novo* PER1 was monitored after existing protein was depleted by 10 hr CHX treatment and washout. Mutant PER1 exhibited accelerated phosphorylation kinetics in gross phosphorylation based on mobility shift (n = 2). **g**, $^{15}N/^{1}H$ HSQC of PER1 FASP peptides corresponding to select indel mutants, with and without CK1$\delta$.
